# Aerolysin nanopore structure revealed at high resolution in lipid environment

**DOI:** 10.1101/2024.08.12.607338

**Authors:** Jana S. Anton, Ioan Iacovache, Juan F. Bada Juarez, Luciano A. Abriata, Louis W. Perrin, Chan Cao, Maria J. Marcaida, Benoit Zuber, Matteo Dal Peraro

## Abstract

Aerolysin is a β-pore-forming toxin produced by most *Aeromonas* bacteria which has attracted large attention in the field of nanopore sensing due to its narrow and charged pore lumen. Structurally similar proteins, belonging to the aerolysin-like family, are present throughout all kingdoms of life, but very few of them have been structurally characterized in a lipid environment. Here we present the first high-resolution atomic cryo-EM structures of aerolysin pre-pore and pore in a membrane-like environment. These structures allow the identification of key interactions, which are relevant for the pore formation and for positioning the pore barrel into the membrane with the anchoring β-turn motif now finally observed. Moreover, we elucidate at high resolution the architecture of key mutations and precisely identify four constriction rings in the pore lumen that are highly relevant for nanopore sensing experiments.

## Introduction

Pore-forming toxins (PFTs) are important virulence factors of many pathogenic bacteria, which are secreted in a soluble form and upon host membrane binding, they oligomerize into transmembrane pores undergoing a complex structural reorganization ^1^. Aerolysin, a major virulence factor of *Aeromonas hydrophila*, is one of the best characterized β-pore-forming toxins ^1–3^. The structure of the soluble form of aerolysin was solved 30 years ago by X-ray crystallography ^4^. The 52 kDa monomer of aerolysin is composed of four domains with different roles in its mode of action (**Figure 1a**): the first two N-terminal domains (domains 1 and 2) are required for receptor binding; the domain 3 forms most of the transmembrane pore; the domain 4, which is the most C-terminal domain, functions as a chaperone and mediates pore formation^5–7^. While the structures of the intermediate oligomers have been solved by cryo-EM at medium resolution (3.5 – 4.5 Å), the structure of the mature aerolysin pore has been so far elusive ^8^. Only a crude model of the pore in detergent micelles has been obtained from a low resolution map (∼8 Å) ^8^.

**Figure 1.**
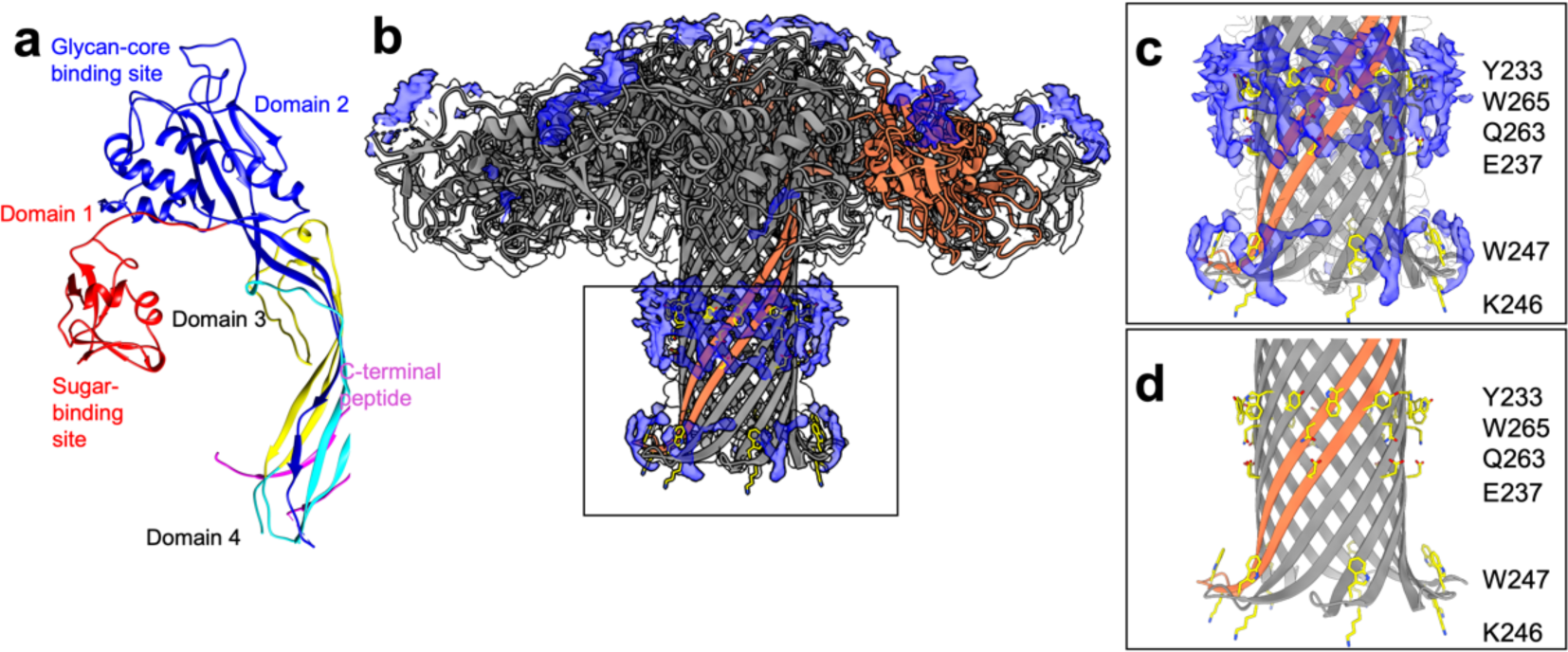
| High resolution structures of wild-type aerolysin in SMALPs. **a** Structure of proaerolysin (PDB ID: 1PRE). Domain 1 is colored in red, domain 2 is colored in dark blue, domains 3 and 4 are colored in turquoise for the part that forms the concentric β-barrel and in yellow for the part that forms the stem β-barrel. The C-terminal peptide is colored in pink. **b** Structure and density map of aerolysin in SMALP. Extra density is highlighted in blue. One protomer is highlighted in orange. **c** Zoom into the lower barrel of aerolysin WT with extra densities observed in the SMALP density map shown in blue. Membrane lining residues are highlighted in yellow. **d** Aerolysin barrel without density map.

Since then, aerolysin has gained particular interest in the field of nanopore sensing due to the unique structural and electrostatic features of its narrow inner pore ^9,10^. In nanopore sensing, the specific nature of an analyte is determined by analyzing its current signature and residence time as it translocates through a pore ^11^. The long, narrow, and charged lumen of aerolysin is characterized by two distinct constrictions narrower than one nanometer. It has enabled the detection and analysis of diverse biomolecules and the discrimination of single amino acids, nucleotides and glycans^12–16^. The introduction of mutations in aerolysin has been shown to increase its sensitivity. For instance, the K238A mutation leads to a large increase in the selectivity and sensitivity of the pore presumably by modifying the size of its constrictions ^13^. Also other mutants, such as K238A-K244A, R220A and T240R, have also shown improved sensing capabilities by modifying one of the two constriction sites of the pore ^16,17^. However, a high-resolution structure of aerolysin is currently lacking, hindering a more precise understanding of its biological function and sensing capabilities.

Here, we present the first high-resolution cryo-EM maps and atomic models of wild-type (WT) in a membrane-like environment and Y221G pre-pore like mutant. In addition, we also solve some key mutants that are regularly used in nanopore experiments, namely K238A and K238A/K244A. These high-resolution structures allow the identification of important interactions required for pore formation and reveal four constriction rings in the pore lumen.

## Results and Discussion

### CryoEM structure of wild-type aerolysin in multiple membrane mimics

The type of membrane mimetic used to solubilize membrane proteins can affect protein structure ^18,19^. To address this issue in the context of β-barrel proteins, we decided to compare the pores obtained in lauryl maltose neopentyl glycol-cholesteryl hemisuccinate (LMNG-CHS), amphipols ^20^ as well as styrene maleic-acid lipid particles (SMALPs)^21^. We obtained near-atomic resolution (2.2 Å) cryo-EM maps of WT aerolysin solubilized in either amphipols or SMALP, as well as in LMNG-CHS (2.6 Å) (**Figure 1b** and **Supplementary Figures 1**). Interestingly, we did not observe any major differences between the density maps and models derived from them (**Supplementary Figure 1**, **Supplementary Table 1 and 2**). This result may seem to contradict previous results that have shown that different solubilization methods impact membrane protein structure in different ways ^19^. However, β-PFTs are highly stable β-barrel structures and all methods of sample preparation that we tested seem equally suited for their structural characterization, a property that can likely translate to all symmetric oligomeric β-barrel proteins. The overall Cα root mean square deviation (RMSD) between the structures in amphipol and SMALP is 0.82 Å for the overall structures and 0.74 Å for the extended β-barrels region (i.e. residues 215-285) (**Supplementary Tables 1 and 2**). Differences between the previously proposed aerolysin pore model and the new high-resolution structures of aerolysin can be observed particularly in the orientation of sidechains and loops. The overall RMSD between the new structures in nanodisc/amphipol and the previous model is ∼3 Å (∼2.5 Å RMSD considering only the extended β-barrel).

The aerolysin pore shows a mushroom-shaped structure with the cap lying parallel to the lipid bilayer as previously reported ^8^. The density for the outer loops of the cap (residue 15-24), which are involved in receptor binding, are less resolved in the cryo-EM maps and a 3D variability analysis on the cryoEM data indicates a large flexibility in this region (**Supplementary Video 1**).

Domains 3 and 4 form the 120 Å long 14-stranded β-barrel, with each protomer contributing two antiparallel β-strands (**Figure 1a**). The transmembrane pore segment has unclear hydropathy features, so its proper position in the membrane is not obvious as for other proteins. However, the new high resolution structures allow a better determination of the interactions of aerolysin with lipids as well as the precise positions of amino acids side chains. The cryoEM map in SMALPs shows an additional density, not observed with amphipol, which allows to partially model some of the lipids surrounding the β-barrel (**Figure 1c,d**, **Supplementary Figure 2**). These lipids allow to pinpoint the position of the nanodisc around the aromatic belt formed by Y233 and W265 (YW belt, **Figure 1c,d**). This belt, a motif commonly observed in β-barrel proteins ^22,23^, anchors and stabilizes the β-barrel in the lipid membrane.

The β-barrel terminates in a partially hydrophobic β-turn motif (amino acids 246-251,**Figure 1c,d**), that is tilted outward and is embedded into the membrane. Together, the seven β-turns form a motif that locks the pore in the membrane, i.e. the rivet structure previously predicted based on site-directed mutagenesis ^6^, later modeled ^8^ and now finally resolved at high resolution. Interestingly, W247, located at the bottom of the β-barrel, points towards the hydrophobic core of the membrane, anchoring the pore in the membrane as previously proposed. The only amino acid located in the rivet which points out of the membrane towards the polar lipid head groups is K246, providing a further anchoring mechanism (**Figure 1c**).

### Structural insights for the pore-forming mechanism

The extensive structural rearrangements that occur when aerolysin transitions from its pre-pore state (captured by the Y221G mutation ^24^) to the mature pore have been previously described ^8,24,25^. This mechanistic model can be refined on the basis of our high resolution cryo-EM structures, as we also solved the structure of the pre-pore Y221G mutant at 1.9 Å resolution (**Figure 2a** and **Supplementary Figure 3**). The overall Cα RMSD with respect to the previous structure at 3.9 Å resolution (PDB ID: 5JZH) is 0.61 Å. Upon pre-pore assembly domains 1 and 2 of adjacent protomers interact through a series of salt bridges and H-bonds that greatly stabilize the cap region (**Supplementary Table 3**). Furthermore, the formation of the concentric β-barrel fold contributes to the stabilization of the pre-pore through the addition of over 10 H-bonds per subunit. Interestingly, five H-bonds (**Figure 2b**) consolidate the lumen of the concentric β-barrel being present both in the pre-pore and pore configuration. The pre-pore stability is further enhanced by the hydrophobic residues at the interface of the two barrels.^8^ Upon membrane insertion, as previously described, the protein twists around two hinge regions flanking the third domain of the protein ^26^. Pore formation retains the tight network of bonds between protomers observed in the pre-pore, while introducing several additional interactions at the folded hinge regions (**Supplementary Table 3)**. Of note, the inter-subunit interaction of H132 and E64 confirms the previously identified critical role of H132 in oligomerization ^26^ (**Figure 2c**).

**Figure 2.**
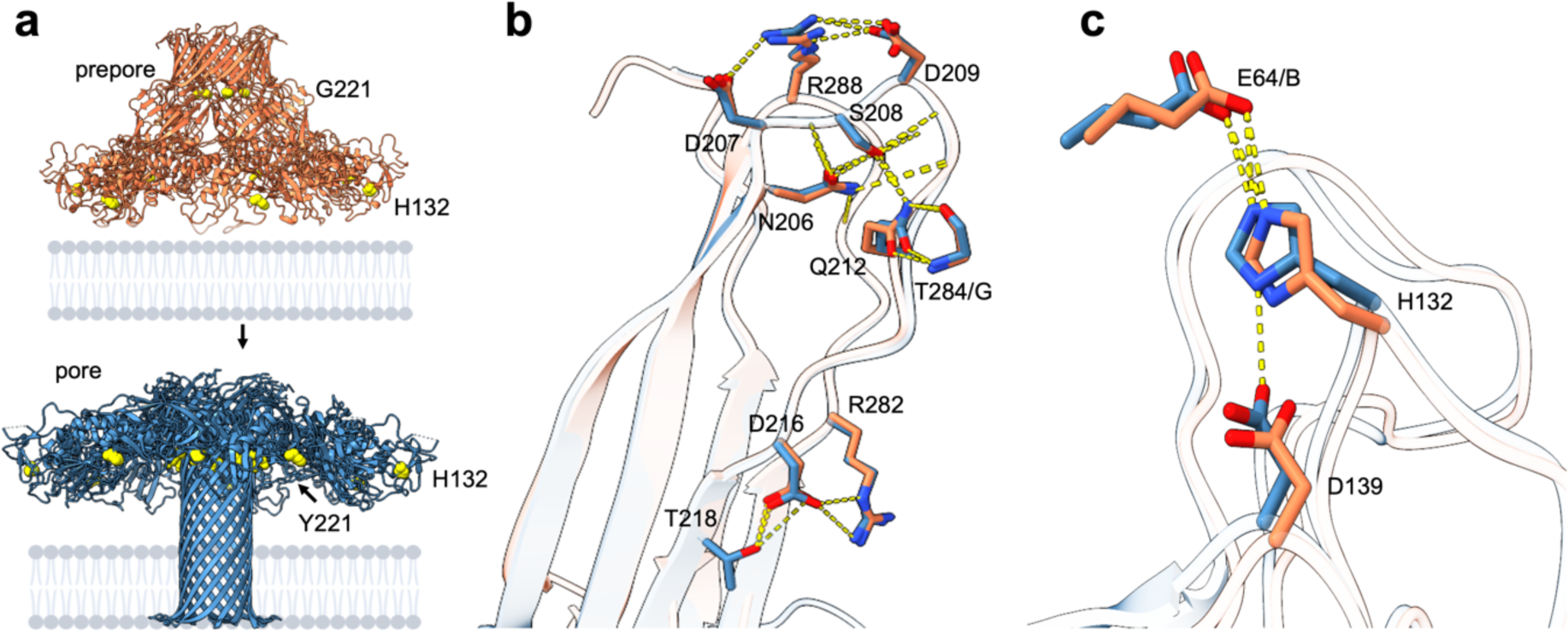
| Structural comparison of Y221G pre-pore and WT pore. **a** Overview of the structural changes from aerolysin pre-pore to pore. The residues H132, Y221 and G221 are highlighted as yellow spheres in the respective structures. **b** Overlay of aerolysin pre-pore and mature pore, zooming in on the concentric β-barrel fold where residues maintain polar interactions between sidechains during the pre-pore to pore transition. These bonds are formed between D207-R288, R288-D209, S208-Q212, Q212-T284 of the neighboring protomer G and R282-D216. **c** Overlay of pre-pore and pore by the cap region highlighting the interaction of residues H132 with D139 and E64 of the neighboring protomer.

### The geometry of the pore lumen and implications for molecular sensing

The geometry of the pore lumen is of particular interest for nanopore sensing applications. Four distinct constriction rings, two at each end of the pore, were identified from the structures (**Figure 3a,b**). The two constrictions on the extracellular end of the barrel are caused by the positively charged amino acids R282 and R220, respectively. R282 forms a salt-bridge with D216 on the same protomer which is involved in further interactions with T218 and S280 (**Figure 3c**). R220 forms polar interactions with D222, which in turn interacts with S276. At the cytoplasmic end of the transmembrane region, the other two constriction sites are defined by K238 and the K242/K244 pair. These lysines form a network of H-bonds and salt-bridges with E258, S256 and E254 on the anti-parallel β-strand of the same or neighboring protomer.

**Figure 3.**
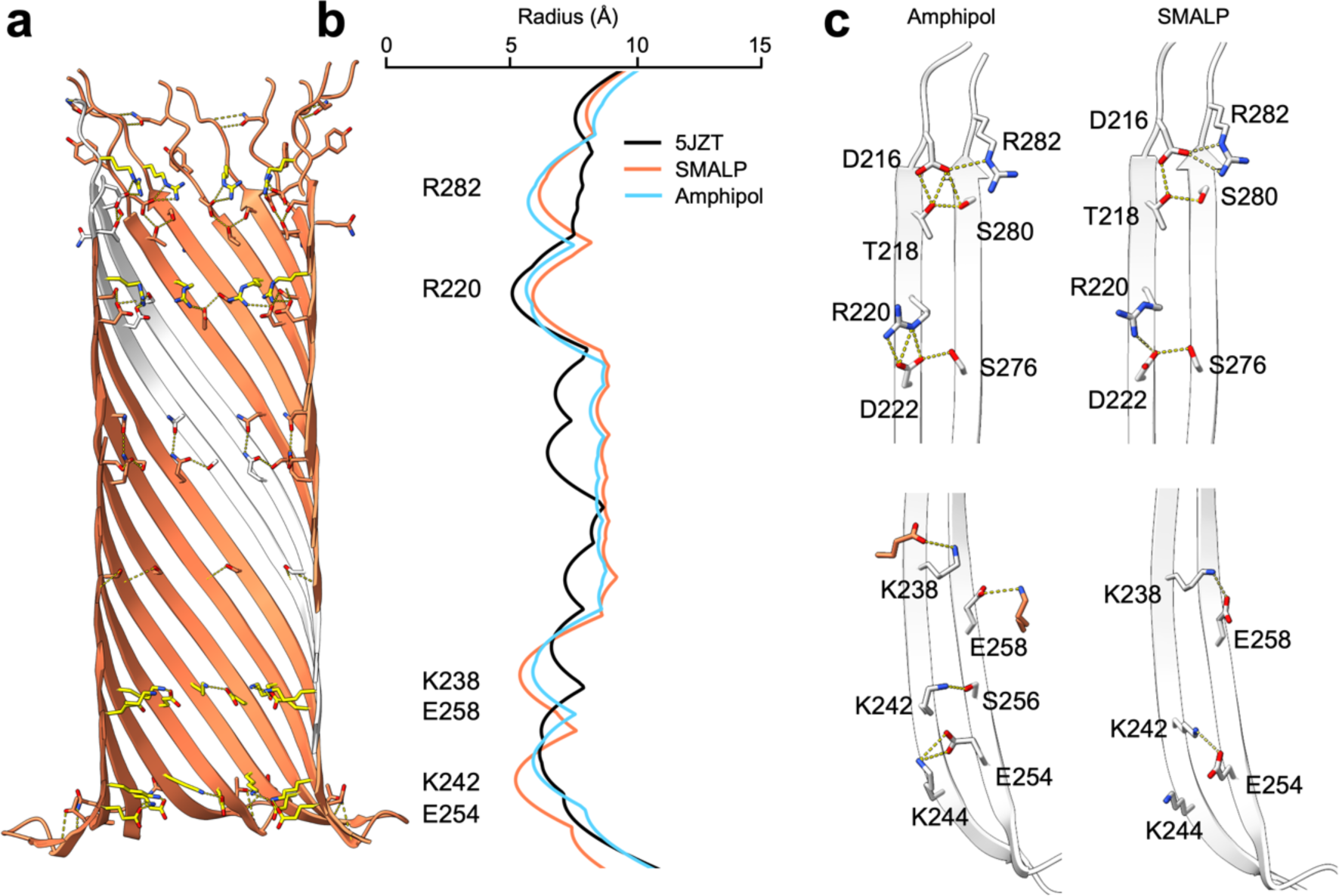
| Origin of aerolysin nanopore sensitivity. **a** Sideview of WT aerolysin lumen in SMALP. Amino acids causing constriction are highlighted in yellow. **b** Traces of the pore radii (Å) of the earlier low-resolution aerolysin structure (5JZT) (black), aerolysin in SMALP (orange) and aerolysin in amphipol (light blue). The radii traces were calculated with HOLE ^27^ from the extracellular to the intracellular end of the pore lumen. **c** Comparison between the network of interactions found in the structures in amphipol and SMALP at the extracellular (top) and intracellular (bottom) ends of the pore.

Interestingly, this interaction network can differ between the WT pore in amphipol and in nanodisc. In amphipol, K242 interacts with S256, while E254 interacts with K244; in the SMALP structure K242 forms a salt-bridge with E254 (**Figure 3c**). These subtle differences in the interaction partners between the two structures suggest that the residues can switch between interaction partners on the same or neighboring protomer, thereby enhancing the stability of the barrel and establishing a robust geometry of the pore lumen. Overall, previously unseen interactions between sidechain and backbone atoms can be observed in both pre-pore and pore conformations of aerolysin structures at high resolution.

The location of the constriction points is in agreement with the sensing spots previously proposed ^12,13^. The central region of the pore lumen exhibits a much wider cavity with a radius of ∼ 8-9 Å, while at the constriction sites the size of the pore radius decreases to ∼ 5 Å. A comparison of the pore radius between the new structures and the previous aerolysin model revealed that only one out of the four constriction rings was clearly captured in the low-resolution model (**Figure 3b**). Similar observations were made when analyzing the water-accessible radius across the pore estimated during MD simulations. In particular, the size of the outermost extracellular constriction formed by R282 was underestimated in the previous aerolysin model. This observation is of particular interest in the field of nanopore engineering where altering constriction size is thought to increase sensitivity ^13,16^.

### CryoEM structure of the K238A aerolysin pore mutant

We have recently demonstrated that the K238A mutation enhances the sensing capabilities and sensitivity due to a longer dwell time of the analyte in the pore ^13,28,29^. To unravel the structural basis for this improved performance, we have solved the structure of the K238A mutant in both amphipol and SMALP, obtaining electron density maps at 2.2 Å. The structures of the wild-type and mutant in SMALP are highly similar, exhibiting an RMSD of 0.92 Å (**Figure 4a**) and comparable pore cavities shown both from the cryoEM structure and in MD simulations (**Figure 4b**). Upon closer examination of the mutant structure, it becomes evident that the K238A mutation induces a widening of the third constriction ring. This is attributed to the substitution of lysines with the smaller alanine residue. In contrast to previous suggestions that the K238A mutation alters the analyte dwell time by decreasing the diameter of the constriction formed by R220 ^13^, these structures reveal that all other constriction rings remain unaffected. Notably, for this mutant, we did observe a narrower water density from MD simulation at R220, which is the same as previous MD results (**Figure 4b)**. Moreover, the structure of the K238A/K244A mutant reveals that also these modifications provide only a local effect (**Supplementary Figure 4**). Interestingly modifying only one of the lysines in the lowest constriction of the pore appears to alter the pore dimension indicating that both K242 and K244 are forming the lowest constriction It has been previously reported that the change in sensitivity was not caused by the impact on the electrostatic potential by the mutation^13^. Consequently, the new structures indicate that the K238A mutation may influence the pore’s rigidity as it disrupts the polar interaction between K238 and E258 within the barrel (**Figure 4c**).

**Figure 4.**
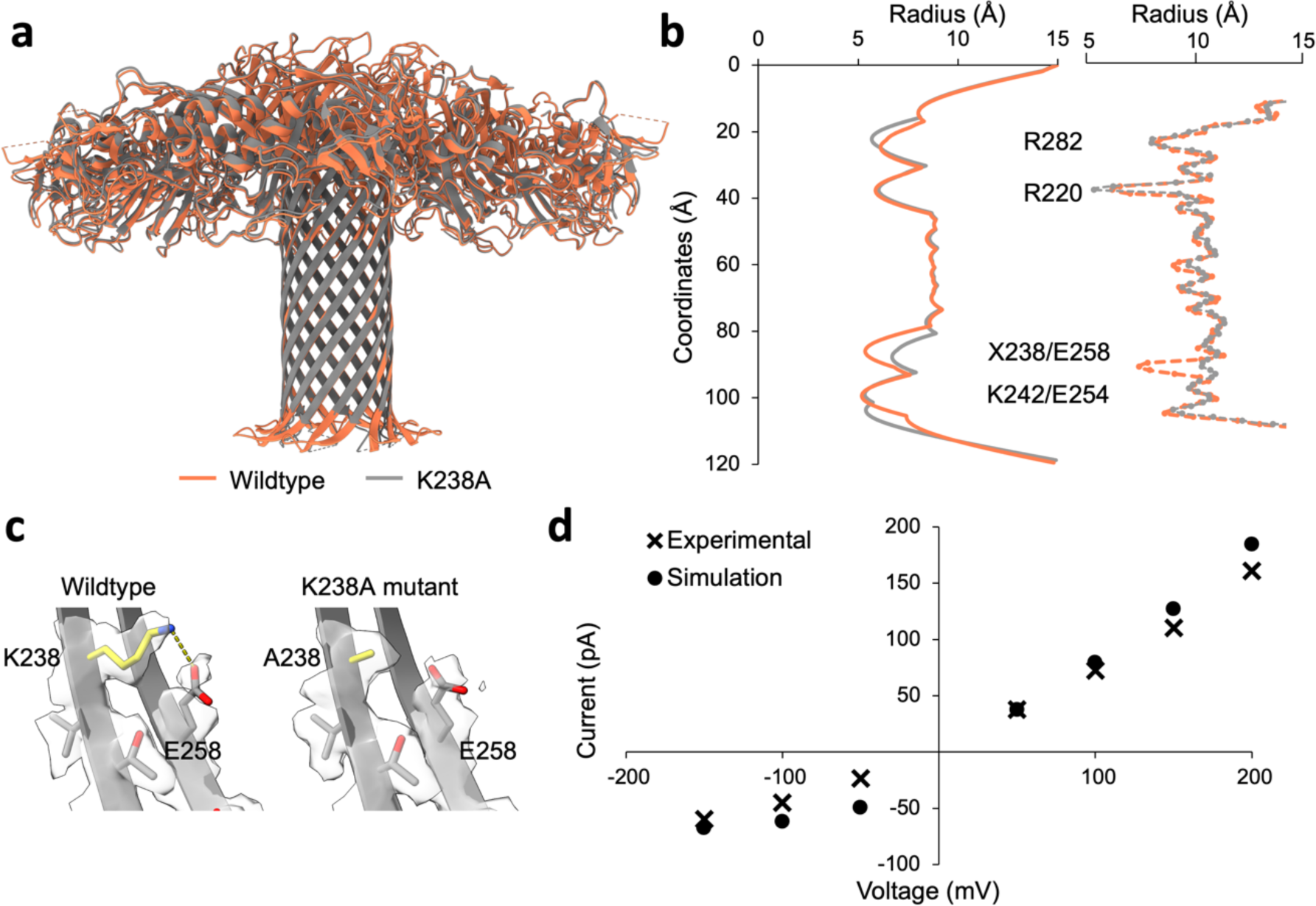
| Structure of the aerolysin K238A mutant used for nanopore sensing. **a** Overlay of WT aerolysin (orange) and aerolysin K238A (gray) in nanodisc. **b** Comparison of the channel radii. On the left side the radii is determined by HOLE ^27^. On the right side the water accessibility radius across the pore is calculated from MD simulations. The amino acid in position 238 depicted by an X is a lysine in the wild-type and alanine in the K238A mutant. **c** Comparison of the polar interactions at the cytoplasmic end of the barrel in WT aerolysin (left) and the K238A mutant (right). The location of the mutation is highlighted in yellow. The density map is shown for residues 238-258. **d** Plot of the current at 1 M KCl as a function of voltage for the K238A aerolysin. Experimental data is shown as crosses^12^, simulation results are shown as circles.

In nanopore sensing experiments the ionic current through the pore is used to determine the nature of the translocating analyte. A comparison of the ionic current response to different voltages revealed that the MD simulations based on the new K238A aerolysin structure solved in SMALPs yielded results that closely aligned with the experimental (**Figure 4d**) ^12,13^.

## Conclusion

In conclusion, we report high resolution structures of wild-type aerolysin along with its K238A and K238A-K244A mutants in SMALPs, amphipols and LMNG. These findings enable us to refine the mechanistic understanding of aerolysin oligomerization and pore formation. We demonstrate that the structural integrity of a β-barrel transmembrane pore is retained across different membrane mimetics, and we reveal for the first time the position of the barrel with respect to the lipid environment. Moreover, our results explain the stability of the aerolysin pore and provide an atomistic view of the interactions conferring the exceptional resilience of the aerolysin pore to denaturation by heat or chaotropic agents ^30^. Of particular interest are the four constriction rings clearly identified in the pore lumen. The accurate structural characterization of these rings and the charge distribution within the pore lumen is of vital importance for the targeted bioengineering of aerolysin pore for tailored nanopore sensing applications. Therefore, our high-resolution structural data hold significant value for the advancement of the fields of nanopore sensing and sequencing ^28,29,31,32^.

### Authors contributions

JSA solved cryoEM structures in nanodisc and analyzed the data, II solved structures in amphipol and LMNG and analyzed the data, JFBJ developed the initial nanodiscs concept and collaborated with JSA to optimize nanodiscs’ formation, LAA and LWP performed and analyzed MD simulations, CC supervised the project and analyzed the data, MJM supervised protein production and structure determination. BZ and MDP designed the project, supervised it and acquired funding. JSA, II, BZ, MDP wrote the paper with input from all authors.

### Accession codes

Aerolysin WT in SMALP: PDB: 9FM6; EMDB: EMD-50549

Aerolysin K238A in SMALP: 9FNP; EMDB: EMD-50601

Aerolysin K238A/K244A in SMALP: 9FNQ; EMDB: EMD-50602

Aerolysin WT in amphipols: 9FML; EMDB: EMD-50562

Aerolysin Y221G: 9FMX; EMDB: EMD-50576

Aerolysin in LMNG:CHS: EMDB: EMD-50578

## Acknowledgements

We thank the laboratory of Prof. G. van der Goot for support in the protein production and for having started the quest for the aerolysin structure. We thank the staff members of the Dubochet Center for Imaging in Lausanne, in particular Dr Emiko Uchikawa and Dr Sergey Nazarov, for their assistance with cryoEM sample preparation and data collection. We thank Prof. Aleksander Antanasijevic and Dr. Yoan Duhoo from EPFL Protein Production and Structure Core Facility for their support in data processing. We acknowledge CSCS, the Swiss National Supercomputing Centre, for access to the HPC resources used to run MD simulations. We thank David Kalbermatter and Marek Kaminek for their support with data acquisition and instrumentation optimization at the Dubochet Center for Imaging in Bern and Microscopy Imaging Cener (MIC) of the University of Bern. We acknowledge the support from the Novartis Foundation, Dementia Research Switzerland (2019-CDA02), the University of Bern Research Foundation (to II), and the Swiss National Science Foundation (grant 200021L_212128 to MDP, and 31003A_179520, 32NE30_185536, CRSII—222809 to BZ, PR00P3_193090 to C.C). Data were acquired on an instrument of the Dubochet Center for Imaging in Lausanne and Bern and supported by the Microscopy Imaging Center (MIC) of the University of Bern.

## Methods

### Aerolysin expression and purification

Wild-type and mutant aerolysin with a C-terminal Hexahistidine tag were expressed using a pET22b vector in BL21 DE3 pLys *E. coli* cells. Cells were grown to an optical density of 0.6 to 0.7 in Luria-Bertani (LB) media shaking at 37°C. Protein expression was induced by adding 1 mM isopropyl β-D-1-thiogalactopyranoside (IPTG) and subsequent growth at 20 °C overnight. Collected cell pellets were resuspended in lysis buffer (20 mM Sodium phosphate pH 7.4, 500 mM NaCl) mixed with cOmplete™ Protease Inhibitor Cocktail (Roche) and subsequently lysed by sonication. After addition of 4 µL of TurboNuclease, the suspension was centrifuged at 20’000x*g* for 30 min at 4°C. The supernatant was applied to an HisTrap HP column (GE Healthcare) previously equilibrated with lysis buffer. Protein was eluted with a gradient over 30 column volumes of elution buffer (20 mM Sodium phosphate pH 7.4, 500 mM NaCl, 500 mM Imidazole). Fractions containing aerolysin were buffer-exchanged into aerolysin buffer (20 mM Tris, pH 8, 150 mM NaCl) using a HiPrep Desalting column (GE Healthcare). Purified protein was frozen in liquid nitrogen and stored at –20 °C until further usage.

### Liposome preparation

For the liposome preparation DOPC and DOPE dissolved in chloroform were mixed in a mol ratio of 2:1 and dried down under a stream of nitrogen. Remaining solvent was removed by incubating the sample in a speedvac for 1h at room temperature. Lipids were resuspended aerolysin buffer (150 mM NaCl, 20 mM Tris, pH 8) to a final concentration of 10 mg/mL. After at least 5 freeze & thawed cycle of the lipid solution, liposomes were formed by extrusion through a 100 nm filter using the mini-extruder from Avanti.

### Preparation of aerolysin in SMALP

For the preparation of aerolysin in SMALP, 9 µM protein, 18 µM liposomes and 0.325 U agarose coupled trypsin were mixed in a final volume of 163 µl and incubated shaking at 150 rpm for 1h at room temperature. After 1h, SMA200 (Cube Biotech) was added in a ratio of 1.5:1 (w/w) polymer to lipid and the mixture was incubated another hour shaking at 150 rpm and room temperature. Subsequently, trypsin was removed by centrifugation at 500*xg* for 10 min at room temperature. The supernatant was used for cryo-EM grid preparation.

### Preparation of aerolysin in A8-35 and LMNG-CHS

For the preparation of Aerolysin in LMNG and A8-35, aerolysin (20 µM) was mixed with 0.05% LMNG-CHS or A8-35 in a final volume of 400 µl. 4 µl agarose-coupled or soluble trypsin (1mg/ml) was added and proteolytic activation was done at 24°C on a rotary shaker for 1h. The sample was dialyzed with a cut-off of 3.5 kDa at room temperature for 2h against 20mM Hepes 100mM NaCl pH 7 prior to vitrification and cryo-EM. In the case of aerolysin Y221G the sample was activated at 8µM and concentrated to 20-30µM after dialysis.

### Cryo-EM grid preparation

For the SMALP datasets, cryo-samples were prepared using a Vitrobot Mark IV (Thermo Fisher Scientific). Grids were frozen at 8°C with a humidity of 95%. Quantifoil R1.2/1.3 on Cu 300 mesh grids coated with 5 nm carbon were glow discharged for 10 s at 10 mA using the GloQube Plus Glow Discharge System from Quorum. 4 µL of sample was applied to the grid and after 1 min incubation the grids were blotted for 4s and quickly plunged into liquid ethane pre-cooled by liquid nitrogen. After vitrification the grids were stored in liquid nitrogen until data collection. For the amphipol and LMNG datasets vitrification was performed at 100% humidity using Quantifoil R1.2/1.3 and R 2/1 with 2nm carbon. Grids were glow discharged on a Baltzer CT010 for 5 s at 10 mA.

### Data collection

The datasets for aerolysin in SMALP were collected at the Dubochet Center for Imaging (Lausanne, CH) using the 300 kV TFS Titan Krios G4 equipped with a Cold-FEG and Falcon 4 detection. The datasets were collected in electron-counting mode (EER). The Falcon IV gain references were measured before starting the data collection. The data collection was performed using the TFS EPU software packages. Movies were recorded at a nominal magnification of 96 000x, corresponding to 0.83 Å/pixel with defocus values ranging from 0.8 to 1.7. The exposure dose was set to 50 e/Å^2^. The amphipol and LMNG datasets were collected at the Dubochet Center for Imaging (Bern, CH) on a 300kV TFS Titan Krios G4 equipped with a Cold-FEG, Falcon 4i and Selectris energy filter with a slit of 20 eV and an average exposure dose set to 40 e/Å^2^. The data was recorded as summarized in the supplements (**Supplementary Table 4**).

### Data processing for aerolysin

All datasets of aerolysin in SMALP were processed in cryoSPARC ^33^. The motion correction was performed on raw stacks without binning using the cryoSPARC Patch motion correction implementation. For all datasets, the initial CTF parameters were estimated using the Patch CTF estimation followed by particle picking using the blob picking tool. After several rounds of 2D classification, an *Ab initio* model was built which was used for template picking. Particles obtained from template picking were again cleaned by several rounds of 2D classification and by building multiple *Ab-initio* models. The selected particles were used for *Ab initio* reconstruction followed by homogenous refinement applying C7 symmetry. The reported resolutions are based on the gold-standard Fourier shell correlation (FSC)= 0.143 criteria^34^ and local-resolution variations were estimated using CryoSPARC.

Amphipol and LMNG datasets were processed in Relion ^35^. In brief, the EER dataset (40 e/Å^2)^ was converted to 40 frames tifs using *relion_convert_to_tiff*. Motion correction ^36^ and CTF-estimation ^37^ were performed followed by particle picking, 2D classification and refinement ^38^. The best particles were polished and CTF refined according to Relion protocols ^39–41^. The reported resolutions are based on the gold-standard FSC=0.143 criterion ^34^. A schematic protocol of the processing for each structure can be found in the supplements (**Supplementary Figures 1a-f**).

### Model building

For model building of the aerolysin in SMALP datasets, the PDB structure of aerolysin (PDB ID: 5JZT) was fit into the cryo-EM map using *Phenix*^42^ and was manually adjusted using *Coot* ^43^ for the K238A mutant. For the other structures, the model of aerolysin wild-type in amphipol was used as an initial model and fit similarly. The final model was generated by iterating between manual model building in *Coot* and relaxed refinement in Rosetta ^44^ and Phenix. In the wt structure in SMALP, the amino acids 15-24 were not built because the density in this area was not well refined. Similarly, residues 10-26 and 246-251 in the K238A mutant in SMALP and residues 13-24 in the K238A-K244A mutant were not built. MolProbidity ^45^ and EMRinger ^46^ were used to validate the final model. The structure was analyzed using UCSF Chimera and UCSF ChimeraX^47,48^. The dimensions of the pores were calculated using the software HOLE ^27^. Figures were prepared in UCSF ChimeraX. Model building of the amphipol structure was performed using ModelAngelo ^49^ followed by Phenix Dock and Rebuild. The best chain was built by iterating between *Coot* and Phenix. For the flexible loops in domain 1 for which the density does not allow accurate modelling the X-ray structure of the aerolysin monomer was used as a guide since the domain was shown to be stable ^30^.

### MD simulation setup

For the all-atom molecular dynamics simulations, aerolysin K238A was embedded into a DPhPC bilayer and the system was prepared (solvated and neutralized) using the CHARMM-GUI software ^50^. The system was equilibrated at 298.15 K for 6 ns. The protonation state of each amino acid was computed for pH 7 and the simulations were performed for 250 ns at 298.15 K at 1 M KCl. The forcefield used was CHARMM36m with the TIP3P water model ^51^. The current at –150 mV, –100 mV, –50 mV, 50 mV, 100 mV, 150 mV and 200 mV were simulated. Experimental data were taken from a previous publication ^12^.

### Water accessibility radius^52^

For the calculation of the hydrodynamic radius, a density map of water and the pore was computed over the box. Based on these density map, with a radius step *dr*, the average density of water and pore in a ring centered on the pore axis with a radius *r*, i.e. the density between *r* and *r+dr* was computed. Once the ratio between the density of water and density of pore reached a threshold value, the last *r* value was retrieved and considered as the hydrodynamic radius. This was then repeated over the pore’s axis to obtain a radius value at each grid point on the pore’s axis.

### Ionic current calculation

The ionic currents were calculated according to the method developed in ^52^. Briefly, the current at time t is computed with the equation hereunder, and then averaged over the trajectory.

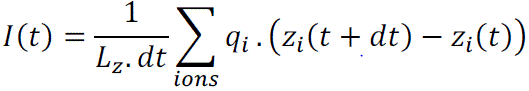

Where *I(t)* is the current, *L_z_* is the box size on the pore’s axis, *dt* is the time step between two frames, *q_i_* is the charge of an ion and *z_i_* is its position on the pore axis.

## Supplementary Information

**Supplementary Table 1:**
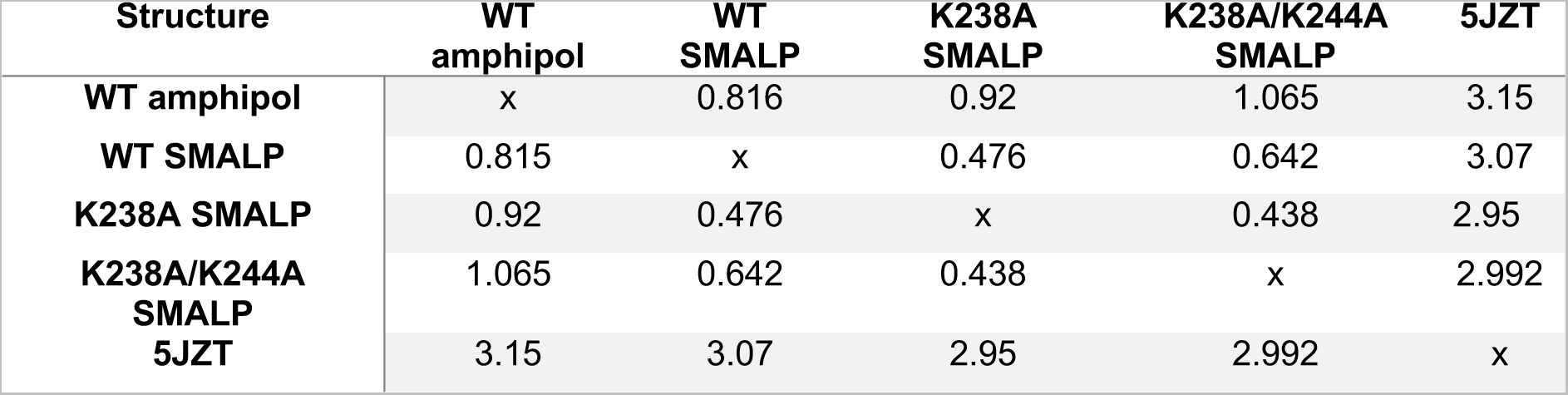
Overview of the overall alpha-carbon RMSD of the aerolysin structures calculated with ChimeraX.

**Supplementary Table 1:**
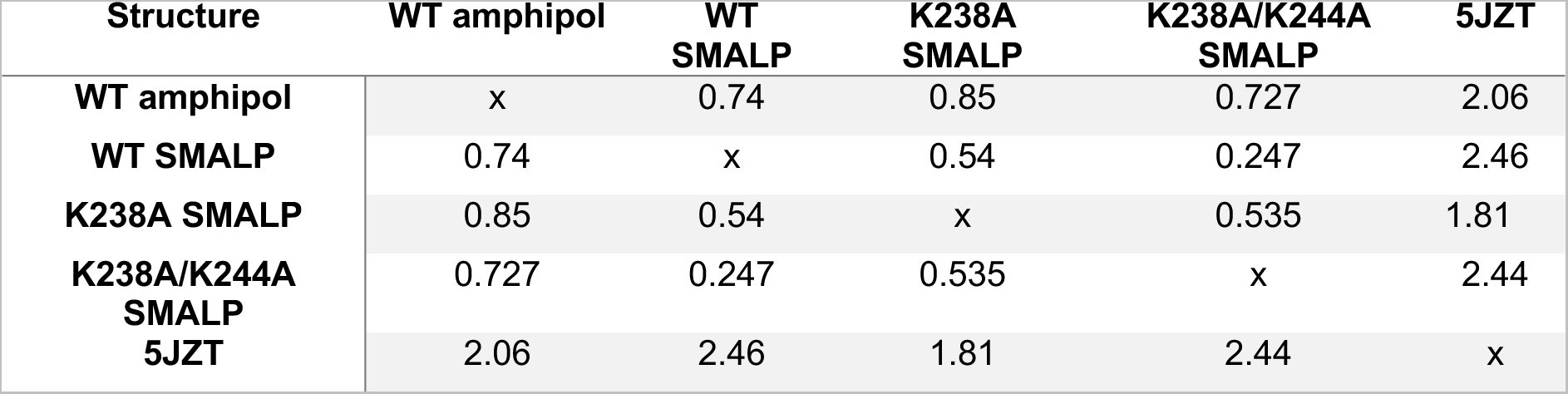
Overview of the alpha-carbon RMSD of the aerolysin barrel (residue 215-285) between the different aerolysin structures. The RMSD was calculated using ChimeraX.

| Structure | WT amphipol | WT SMALP | K238A SMALP | K238A/K244A SMALP | 5JZT |
| --- | --- | --- | --- | --- | --- |
| WT amphipol | x | 0.74 | 0.85 | 0.727 | 2.06 |
| WT SMALP | 0.74 | x | 0.54 | 0.247 | 2.46 |
| K238A SMALP | 0.85 | 0.54 | x | 0.535 | 1.81 |
| K238A/K244A SMALP | 0.727 | 0.247 | 0.535 | x | 2.44 |
| 5JZT | 2.06 | 2.46 | 1.81 | 2.44 | x |

**Supplementary Table 2:** Overview of the hydrogen bonds formed between sidechains in aerolysin wt and amphipol and SMALP and in the Y221G pre-pore mutant. Interactions are only shown for chain A /X represents the amino acid from protomer x. Analysis was performed using ChimeraX.

| WT amphipol |  |  | WT SMALP |  |  | Y221G mutant |  |  |
| --- | --- | --- | --- | --- | --- | --- | --- | --- |
| Donor | Acceptor | Distance (Å) | Donor | Acceptor | Distance (Å) | Donor | Acceptor | Distance (Å) |
| /A TYR 5 | /A TYR 76 | 3.2 | A TYR 5 | /A TYR 76 | 3.2 | /A GLN 8 | /A ASP 7 | 2.8 |
| /A GLN 8 | /A TYR 76 | 3.5 | /A GLN 8 | /A ASP 7 | 3.0 | /A ARG 10 | /A ASP 7 | 2.4 |
| /A GLN 8 | /A VAL 81 | 2.9 | /A ASN 27 | /A GLU 30 | 3.0 | /A ARG 10 | /A LEU 9 | 2.7 |
| /A ARG 10 | /A ASP 7 | 2.3 | /A ARG 28 | /G MET 140 | 3.0 | /A ARG 10 | /A ASP 7 | 3.0 |
| /A ARG 10 | /A LEU 9 | 2.5 | /A ARG 28 | /G ASP 139 | 2.6 | /A ASN 27 | /A GLU 30 | 3.4 |
| /A ARG 10 | /A ASP 7 | 3.0 | /A ARG 28 | /G ASP 141 | 3.3 | /A ARG 28 | /B MET 140 | 2.7 |
| /A ARG 24 | /A PRO 25 | 3.3 | /A GLU 29 | /A GLU 29 | 2.6 | /A ARG 28 | /B ASP 139 | 2.4 |
| /A ARG 28 | /B MET 140 | 3.5 | /A GLN 32 | /G VAL 142 | 3.1 | /A ARG 28 | /B MET 140 | 3.3 |
| /A ARG 28 | /B MET 140 | 3.5 | /A SER 33 | /A GLU 30 | 3.0 | /A ARG 28 | /B ASP 141 | 2.9 |
| /A GLU 30 | /A ASN 27 | 2.8 | /A LYS 35 | /G ASP 156 | 3.4 | /A GLU 30 | /A ASN 27 | 2.5 |
| /A GLN 32 | /B VAL 142 | 3.4 | /A GLY 43 | /A GLN 46 | 2.7 | /A GLN 32 | /B VAL 142 | 3.1 |
| /A SER 33 | /A GLU 30 | 2.6 | /A SER 48 | /A ASN 72 | 2.6 | /A LYS 35 | /A GLY 63 | 2.9 |
| /A LYS 35 | /A GLY 63 | 3.4 | /A TYR 61 | /A TRP 45 | 2.8 | /A GLY 43 | /A GLN 46 | 2.5 |
| /A GLY 43 | /A GLN 46 | 2.6 | /A ASN 72 | /A ILE 47 | 3.1 | /A GLU 84 | /A GLU 84 | 2.7 |
| /A GLN 46 | /A SER 48 | 3.4 | /A ASN 72 | /A SER 71 | 3.2 | /A VAL 99 | /A ASP 97 | 2.7 |
| /A ASN 72 | /A ILE 47 | 3.0 | /A CYS 75 | /A THR 73 | 3.6 | /A GLN 102 | /A TYR 177 | 3.1 |
| /A THR 78 | /A LYS 22 | 2.7 | /A GLU 98 | /A GLU 98 | 2.8 | /A HIS 107 | /A TRP 103 | 3.0 |
| /A ILE 85 | /A GLU 84 | 3.1 | /A ASP 100 | /A ASP 97 | 3.0 | /A ASN 111 | /A ASP 108 | 3.1 |
| /A GLU 98 | /A GLU 98 | 3.0 | /A ARG 104 | /B GLU 2 | 3.0 | /A ASN 111 | /A ASP 108 | 2.7 |
| /A VAL 99 | /A ASP 97 | 3.4 | /A ASN 111 | /A ASP 108 | 3.1 | /A TRP 127 | /A TYR 162 | 3.0 |
| /A ASP 100 | /A ASP 97 | 2.8 | /A ASN 111 | /A ASP 108 | 3.0 | /A ASN 131 | /A ASN 155 | 3.4 |
| /A GLN 102 | /A TYR 177 | 3.4 | /A LYS 114 | /A VAL 136 | 2.7 | /A ASN 131 | /A ASP 156 | 3.0 |
| /A ARG 104 | /G GLU 2 | 3.4 | /A LYS 114 | /A GLU 138 | 2.9 | /A HIS 132 | /A ASP 139 | 2.9 |
| /A HIS 107 | /A TRP 103 | 2.9 | /A THR 116 | /A PHE 112 | 2.8 | /A HIS 132 | /A ASP 139 | 2.8 |
| /A ASN 111 | /A ASP 108 | 3.4 | /A SER 117 | /A ILE 113 | 2.7 | /A HIS 132 | /G GLU 64 | 0.0 |
| /A ASN 111 | /A ASP 108 | 3.4 | /A TRP 127 | /A TYR 162 | 3.1 | /A SER 133 | /A GLU 138 | 2.7 |
| /A ASN 111 | /A ASP 108 | 3.0 | /A ASN 131 | /A ASN 155 | 2.9 | /A TYR 135 | /A SER 133 | 3.2 |
| /A LYS 114 | /A VAL 136 | 2.5 | /A ASN 131 | /A GLY 157 | 3.1 | /A VAL 142 | /G GLN 32 | 3.1 |
| /A THR 116 | /A PHE 112 | 3.1 | /A HIS 132 | /A ASP 139 | 2.8 | /A ARG 144 | /G GLU 367 | 3.0 |
| /A SER 117 | /A ILE 113 | 2.8 | /A HIS 132 | /A ASP 139 | 3.0 | /A ARG 144 | /A GLY 148 | 3.5 |
| /A TYR 125 | /A LEU 318 | 3.4 | /A HIS 132 | /B GLU 64 | 3.1 | /A TRP 149 | /A GLN 102 | 3.1 |
| /A ASN 131 | /A ASN 155 | 3.4 | /A GLN 134 | /A GLN 134 | 2.6 | /A ARG 163 | /A ASP 166 | 2.9 |
| /A HIS 132 | /A ASP 139 | 3.3 | /A TYR 135 | /A SER 133 | 3.1 | /A ARG 163 | /A ARG 336 | 3.3 |
| /A HIS 132 | /A ASP 139 | 2.7 | /A VAL 142 | /B GLN 32 | 2.9 | /A ARG 163 | /A ASP 166 | 3.2 |
| /A HIS 132 | /G GLU 64 | 2.8 | /A THR 143 | /A ARG 144 | 3.6 | /A LYS 167 | /A ASN 131 | 2.7 |
| /A VAL 142 | /G GLN 32 | 3.2 | /A ARG 144 | /B GLU 367 | 3.1 | /A LYS 167 | /A ASN 155 | 2.8 |
| /A THR 143 | /A VAL 150 | 3.5 | /A ARG 144 | /A GLN 102 | 3.2 | /A ASN 174 | /A ASP 315 | 3.0 |
| /A ARG 144 | /G GLU 367 | 3.2 | /A TRP 149 | /A GLN 102 | 2.9 | /A TYR 177 | /A GLU 98 | 2.9 |
| /A ARG 144 | /A GLN 102 | 3.3 | /A ASN 154 | /A ASP 141 | 2.9 | /A ASP 182 | /A ASP 180 | 3.2 |
| /A TRP 149 | /A GLN 102 | 3.1 | /A ASN 154 | /A ASP 156 | 2.9 | /A HIS 186 | /A GLY 187 | 2.8 |
| /A ASN 154 | /A ASP 141 | 2.6 | /A ASP 156 | /A ASN 154 | 3.0 | /A SER 192 | /A ASP 193 | 3.5 |
| /A ASN 154 | /A ASP 156 | 3.3 | /A ARG 163 | /A ASP 166 | 3.0 | /A ASN 206 | /A THR 210 | 2.9 |
| /A ASP 156 | /A ASN 154 | 3.3 | /A ARG 163 | /A ASP 166 | 3.1 | /A ASN 206 | /A VAL 285 | 2.8 |
| /A LYS 167 | /A GLY 129 | 3.0 | /A LYS 167 | /A GLY 129 | 2.9 | /A SER 208 | /A ASN 206 | 2.9 |
| /A LYS 167 | /A ASN 131 | 3.1 | /A LYS 167 | /A ASN 131 | 2.9 | /A SER 208 | /A ASN 206 | 2.8 |
| /A THR 168 | /A ALA 126 | 2.9 | /A THR 168 | /A ALA 126 | 2.6 | /A THR 210 | /A SER 208 | 3.2 |
| /A ASN 174 | /A ASP 315 | 3.5 | /A ASN 174 | /A ASP 315 | 2.9 | /A GLN 212 | /A SER 208 | 3.3 |
| /A SER 183 | /A ASP 180 | 3.0 | /A SER 183 | /A ASP 180 | 2.9 | /A GLN 212 | /G THR 284 | 3.4 |
| /A SER 183 | /A LYS 309 | 3.4 | /A SER 183 | /A LYS 309 | 3.5 | /A TYR 215 | /A VAL 205 | 2.6 |
| /A LYS 185 | /A GLU 307 | 2.8 | /A HIS 186 | /A GLY 187 | 3.3 | /A THR 225 | /A THR 273 | 3.0 |
| /A HIS 186 | /A GLY 187 | 2.9 | /A HIS 186 | /G GLU 415 | 2.9 | /A LYS 229 | /A TRP 265 | 2.6 |
| /A HIS 186 | /B GLU 415 | 2.9 | /A THR 190 | /A SER 303 | 3.5 | /A LYS 229 | /A GLN 268 | 2.5 |
| /A HIS 186 | /B GLU 415 | 3.1 | /A ARG 194 | /A GLN 191 | 2.8 | /A ASN 231 | /A ALA 266 | 3.5 |
| /A THR 190 | /A SER 303 | 3.4 | /A GLN 195 | /A ASP 193 | 3.1 | /A ASN 231 | /A ASN 269 | 2.8 |
| /A SER 192 | /A ASP 193 | 3.0 | /A LYS 198 | /B SER 408 | 3.3 | /A LYS 242 | /A ASN 178 | 3.2 |
| /A ARG 194 | /A GLN 191 | 2.6 | /A THR 199 | /B GLN 409 | 2.8 | /A TRP 247 | /A TYR 304 | 2.8 |
| /A ARG 194 | /A GLN 191 | 3.1 | /A THR 199 | /B GLN 409 | 3.3 | /A ALA 260 | /A GLN 268 | 3.0 |
| /A THR 199 | /G GLN 409 | 3.1 | /A ASN 206 | /A THR 210 | 3.0 | /A TRP 265 | /A TYR 306 | 3.1 |
| /A THR 199 | /A ILE 295 | 3.0 | /A ASN 206 | /A VAL 285 | 3.1 | /A SER 267 | /A SER 264 | 2.6 |
| /A ASN 206 | /A THR 210 | 2.8 | /A SER 208 | /A ASN 206 | 2.9 | /A SER 267 | /A SER 264 | 3.3 |
| /A ASN 206 | /A VAL 285 | 3.1 | /A SER 208 | /A ASN 206 | 2.8 | /A GLN 268 | /A ALA 260 | 2.9 |
| /A ASN 206 | /A PRO 286 | 3.1 | /A THR 210 | /A SER 208 | 3.3 | /A GLN 268 | /A GLN 263 | 3.3 |
| /A SER 208 | /A ASN 206 | 3.0 | /A THR 210 | /A PRO 211 | 3.4 | /A ASN 269 | /A ALA 266 | 2.7 |
| /A SER 208 | /A ASN 206 | 2.5 | /A GLN 212 | /A SER 208 | 3.3 | /A GLN 279 | /A GLU 296 | 3.0 |
| /A SER 208 | /A THR 210 | 3.2 | /A GLN 212 | /B THR 284 | 3.3 | /A ARG 282 | /A ASP 216 | 2.5 |
| /A THR 210 | /A PRO 211 | 3.5 | /A TYR 215 | /A VAL 205 | 2.8 | /A THR 284 | /B GLN 212 | 3.1 |
| /A GLN 212 | /A SER 208 | 3.0 | /A THR 218 | /A ASP 216 | 2.8 | /A ARG 288 | /A ASP 207 | 3.1 |
| /A GLN 212 | /G THR 284 | 3.3 | /A THR 218 | /A SER 280 | 2.7 | /A ARG 288 | /A ASP 209 | 3.5 |
| /A SER 213 | /A THR 284 | 3.2 | /A ARG 220 | /A ASP 222 | 3.2 | /A ARG 288 | /A ASP 209 | 3.1 |
| /A TYR 215 | /A VAL 205 | 2.9 | /A SER 228 | /B GLN 268 | 2.7 | /A SER 289 | /A PRO 286 | 2.6 |
| /A THR 218 | /A ASP 216 | 2.9 | /A LYS 238 | /A GLU 258 | 2.8 | /A ASP 301 | /A SER 192 | 2.8 |
| /A THR 218 | /A ASP 216 | 3.0 | /A LYS 242 | /A GLU 254 | 2.9 | /A TYR 306 | /A THR 230 | 2.9 |
| /A THR 218 | /A SER 280 | 2.3 | /A ASN 243 | /B GLU 252 | 2.9 | /A LYS 309 | /A SER 183 | 2.9 |
| /A ARG 220 | /A ASP 222 | 3.2 | /A ASN 243 | /B THR 253 | 2.7 | /A LYS 309 | /A GLU 307 | 3.0 |
| /A ARG 220 | /A ASP 222 | 3.0 | /A THR 253 | /A GLY 251 | 3.4 | /A ARG 323 | /A HIS 121 | 3.0 |
| /A ARG 220 | /A ASP 222 | 2.9 | /A SER 264 | /A TYR 233 | 3.5 | /A ARG 323 | /A TYR 125 | 3.3 |
| /A SER 228 | /A ASN 269 | 3.4 | /A GLN 268 | /G ASN 226 | 3.0 | /A ASN 327 | /A ARG 323 | 3.2 |
| /A SER 236 | /A ALA 261 | 3.3 | /A SER 276 | /A ASP 222 | 2.7 | /A ASN 327 | /A PRO 333 | 2.8 |
| /A LYS 238 | /G GLU 258 | 3.4 | /A GLN 279 | /A GLU 296 | 3.4 | /A TRP 329 | /A ASN 327 | 3.2 |
| /A LYS 242 | /A SER 256 | 2.7 | /A SER 280 | /A THR 218 | 2.7 | /A HIS 332 | /A ASN 327 | 3.0 |
| /A LYS 244 | /A GLU 254 | 3.0 | /A ARG 282 | /A ASP 216 | 2.7 | /A ASN 340 | /A SER 319 | 2.8 |
| /A LYS 244 | /A GLU 254 | 3.3 | /A ARG 282 | /A ASP 216 | 3.1 | /A LYS 351 | /A TYR 357 | 3.5 |
| /A THR 253 | /A GLY 251 | 3.2 | /A THR 284 | /G GLN 212 | 2.9 | /A SER 353 | /A ASP 350 | 3.0 |
| /A THR 253 | /B ASN 243 | 3.4 | /A ARG 288 | /A ASP 209 | 3.0 | /A ARG 356 | /A ILE 85 | 3.4 |
| /A GLN 263 | /A GLU 237 | 3.2 | /A ARG 288 | /A ASP 207 | 3.0 | /A ARG 356 | /A ILE 85 | 3.5 |
| /A GLN 268 | /A SER 228 | 2.5 | /A ARG 288 | /A ASP 209 | 3.0 | /A TYR 357 | /A SER 354 | 2.9 |
| /A GLN 268 | /B ASN 226 | 3.1 | /A SER 289 | /A PRO 286 | 2.7 | /A GLN 358 | /A LYS 351 | 3.1 |
| /A SER 276 | /A ASP 222 | 2.4 | /A LYS 290 | /C PRO 181 | 3.0 | /A GLN 358 | /A ALA 352 | 2.9 |
| /A GLN 279 | /A GLU 296 | 3.3 | /A LYS 294 | /A GLU 296 | 2.9 | /A LYS 361 | /B ASP 97 | 3.2 |
| /A SER 280 | /A ASP 216 | 3.3 | /A TYR 298 | /A GLU 296 | 2.9 | /A ARG 362 | /A TRP 359 | 2.9 |
| /A SER 280 | /A THR 218 | 2.3 | /A SER 303 | /A PHE 404 | 3.3 | /A TRP 373 | /A GLN 358 | 2.9 |
| /A ARG 282 | /A ASP 216 | 3.1 | /A SER 303 | /B TYR 348 | 2.8 | /A ASN 374 | /A ARG 362 | 2.9 |
| /A THR 284 | /B GLN 212 | 3.1 | /A TYR 304 | /G GLU 296 | 2.8 | /A TRP 375 | /A ASP 372 | 3.0 |
| /A THR 284 | /A SER 213 | 3.2 | /A TYR 306 | /A GLU 98 | 2.8 | /A ASN 380 | /A THR 376 | 3.4 |
| /A SER 289 | /A PRO 286 | 2.3 | /A LYS 309 | /A SER 183 | 3.1 | /A GLN 386 | /A GLU 84 | 3.4 |
| /A SER 289 | /A ALA 287 | 3.4 | /A SER 313 | /A ALA 176 | 3.5 | /A ASN 388 | /A PRO 115 | 3.0 |
| /A LYS 290 | /F PRO 181 | 2.5 | /A THR 317 | /A THR 342 | 2.7 | /A ARG 394 | /A SER 89 | 3.0 |
| /A LYS 299 | /A SER 408 | 2.8 | /A SER 319 | /A ASN 340 | 2.7 | /A ARG 397 | /A ASP 92 | 2.9 |
| /A LYS 299 | /A GLN 409 | 3.4 | /A ASN 327 | /A ARG 323 | 3.4 | /A ARG 397 | /A ASP 92 | 2.9 |
| /A ASP 301 | /A ASP 193 | 3.0 | /A ASN 327 | /A PRO 333 | 3.2 | /A THR 401 | /A ASP 95 | 3.2 |
| /A SER 303 | /A SER 405 | 2.9 | /A TRP 329 | /A ASN 327 | 3.0 | /A THR 401 | /A ASP 95 | 2.9 |
| /A SER 303 | /G TYR 348 | 3.1 | /A HIS 332 | /A ASN 327 | 3.0 | /A ASP 403 | /A THR 230 | 3.0 |
| /A TYR 304 | /B GLU 296 | 3.1 | /A THR 342 | /A THR 317 | 2.7 | /A ASN 413 | /A GLU 296 | 3.0 |
| /A TYR 306 | /A GLU 98 | 2.9 | /A LYS 349 | /G SER 192 | 3.0 | /A ASN 413 | /A GLU 415 | 2.8 |
| /A LYS 309 | /A SER 183 | 2.9 | /A LYS 349 | /G ASP 301 | 3.0 |  |  |  |
| /A SER 313 | /A ALA 176 | 3.4 | /A LYS 351 | /A ASP 350 | 2.6 |  |  |  |
| /A THR 317 | /A THR 342 | 3.1 | /A ARG 356 | /A ILE 85 | 3.2 |  |  |  |
| /A SER 319 | /A ASN 340 | 2.3 | /A TYR 357 | /A SER 354 | 3.2 |  |  |  |
| /A ARG 323 | /A HIS 121 | 3.3 | /A TYR 357 | /A GLU 367 | 2.8 |  |  |  |
| /A ARG 323 | /A TYR 125 | 2.5 | /A GLN 358 | /A LYS 351 | 3.1 |  |  |  |
| /A ARG 323 | /A HIS 121 | 3.3 | /A GLN 358 | /A ALA 352 | 3.1 |  |  |  |
| /A ASN 327 | /A ARG 323 | 3.2 | /A TRP 359 | /A GLY 83 | 2.8 |  |  |  |
| /A ASN 327 | /A PRO 333 | 3.0 | /A LYS 361 | /G ASP 97 | 2.8 |  |  |  |
| /A TRP 329 | /A ASN 327 | 3.2 | /A ARG 362 | /A TRP 359 | 3.2 |  |  |  |
| /A HIS 332 | /A ASN 327 | 2.9 | /A ARG 362 | /A THR 82 | 2.8 |  |  |  |
| /A ASN 338 | /A GLY 320 | 3.0 | /A TYR 363 | /G ASP 100 | 2.6 |  |  |  |
| /A THR 342 | /A THR 317 | 3.1 | /A TRP 373 | /A GLN 358 | 3.2 |  |  |  |
| /A LYS 351 | /A TYR 357 | 2.7 | /A ASN 374 | /A ASP 372 | 2.8 |  |  |  |
| /A SER 353 | /A ASP 350 | 2.9 | /A ASN 374 | /A ARG 362 | 3.1 |  |  |  |
| /A SER 353 | /A ASP 350 | 3.4 | /A TRP 375 | /A ASP 372 | 3.0 |  |  |  |
| /A SER 353 | /A ASP 350 | 3.1 | /A THR 376 | /A TRP 373 | 2.9 |  |  |  |
| /A ARG 356 | /A ILE 85 | 3.4 | /A ASN 380 | /A THR 376 | 3.2 |  |  |  |
| /A TYR 357 | /A GLU 367 | 2.9 | /A THR 384 | /A ASN 380 | 2.8 |  |  |  |
| /A GLN 358 | /A LYS 351 | 3.1 | /A GLN 386 | /A GLU 84 | 2.9 |  |  |  |
| /A GLN 358 | /A ALA 352 | 3.2 | /A ARG 394 | /A SER 89 | 2.8 |  |  |  |
| /A TRP<br>359 | /A GLY 83 | 2.6 | /A ARG<br>397 | /A ASP<br>311 | 3.0 |  |  |  |
| /A LYS<br>361 | /B ASP 97 | 3.0 | /A ARG<br>397 | /A ASP<br>180 | 3.4 |  |  |  |
| /A LYS<br>361 | /B ASP<br>100 | 3.2 | /A ARG<br>397 | /A ASP<br>180 | 2.7 |  |  |  |
| /A ARG<br>362 | /A TRP<br>359 | 2.4 | /A GLY<br>399 | /A ASP 95 | 2.9 |  |  |  |
| /A TYR<br>363 | /B ASP<br>100 | 2.4 | /A SER<br>405 | /B LYS<br>349 | 2.8 |  |  |  |
| /A TRP<br>373 | /A GLN<br>358 | 2.9 | /A SER<br>408 | /A ASP<br>301 | 3.0 |  |  |  |
| /A ASN<br>374 | /A ARG<br>362 | 3.0 | /A GLN<br>409 | /A GLU<br>407 | 2.8 |  |  |  |
| /A TRP<br>375 | /A ASP<br>372 | 3.4 |  |  |  |  |  |  |
| /A THR<br>376 | /A TRP<br>373 | 2.7 |  |  |  |  |  |  |
| /A ASN<br>380 | /A THR<br>376 | 3.2 |  |  |  |  |  |  |
| /A THR<br>384 | /A ASN<br>380 | 3.5 |  |  |  |  |  |  |
| /A GLN<br>386 | /A GLU 84 | 2.9 |  |  |  |  |  |  |
| /A ASN<br>388 | /A PRO<br>115 | 3.0 |  |  |  |  |  |  |
| /A ARG<br>394 | /A SER 89 | 2.5 |  |  |  |  |  |  |
| /A ARG<br>394 | /A SER 89 | 3.2 |  |  |  |  |  |  |
| /A ARG<br>394 | /A SER 89 | 3.1 |  |  |  |  |  |  |
| /A ARG<br>397 | /A ASP<br>180 | 3.5 |  |  |  |  |  |  |
| /A ARG<br>397 | /A ASP<br>180 | 3.5 |  |  |  |  |  |  |
| /A ARG<br>397 | /A ASP<br>311 | 3.5 |  |  |  |  |  |  |
| /A ARG<br>397 | /A ASP<br>180 | 2.8 |  |  |  |  |  |  |
| /A SER<br>405 | /A SER<br>303 | 2.9 |  |  |  |  |  |  |
| /A SER<br>405 | /G TYR<br>348 | 3.5 |  |  |  |  |  |  |
| /A SER<br>405 | /G LYS<br>349 | 3.1 |  |  |  |  |  |  |
| /A SER<br>408 | /A ASP<br>301 | 2.9 |  |  |  |  |  |  |
| /A GLN<br>409 | /A GLU<br>407 | 2.7 |  |  |  |  |  |  |

**Supplementary Figure 1:**
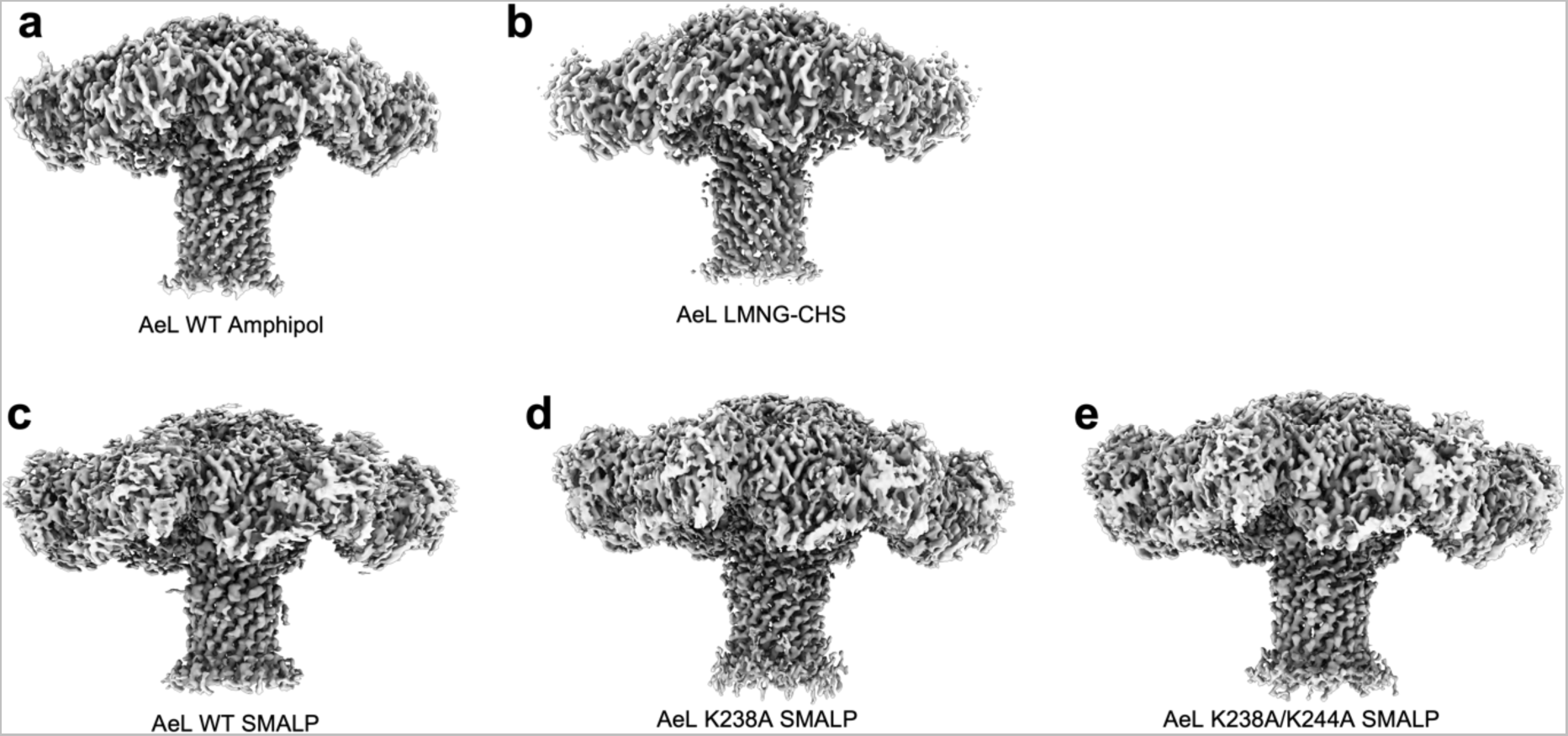
Overview of the density maps generated from cryoEM.

**Supplementary Figure 1a:**
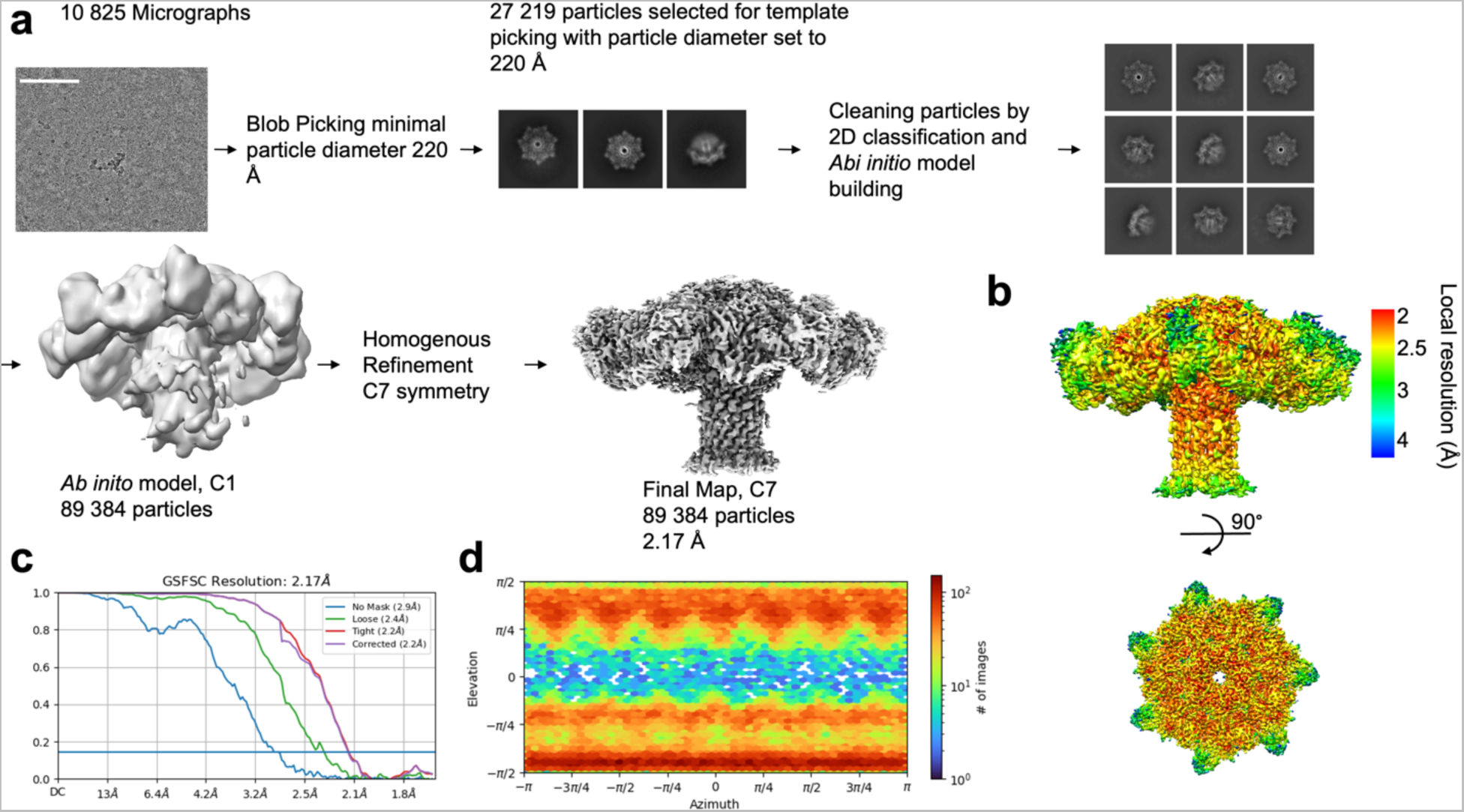
Cryo-EM processing of aerolysin WT in SMALP. **a** Flow chart of the data processing in cryoSPARC. Scale bar represent 100 nm. **b** Final 3D reconstruction of aerolysin wild-type in SMALP colored by resolution. **c** Golden-standard Fourier shell correlation curves for the 3D reconstruction. **d** Angular distribution of the particles included in the final reconstruction.

**Supplementary Figure 1b:**
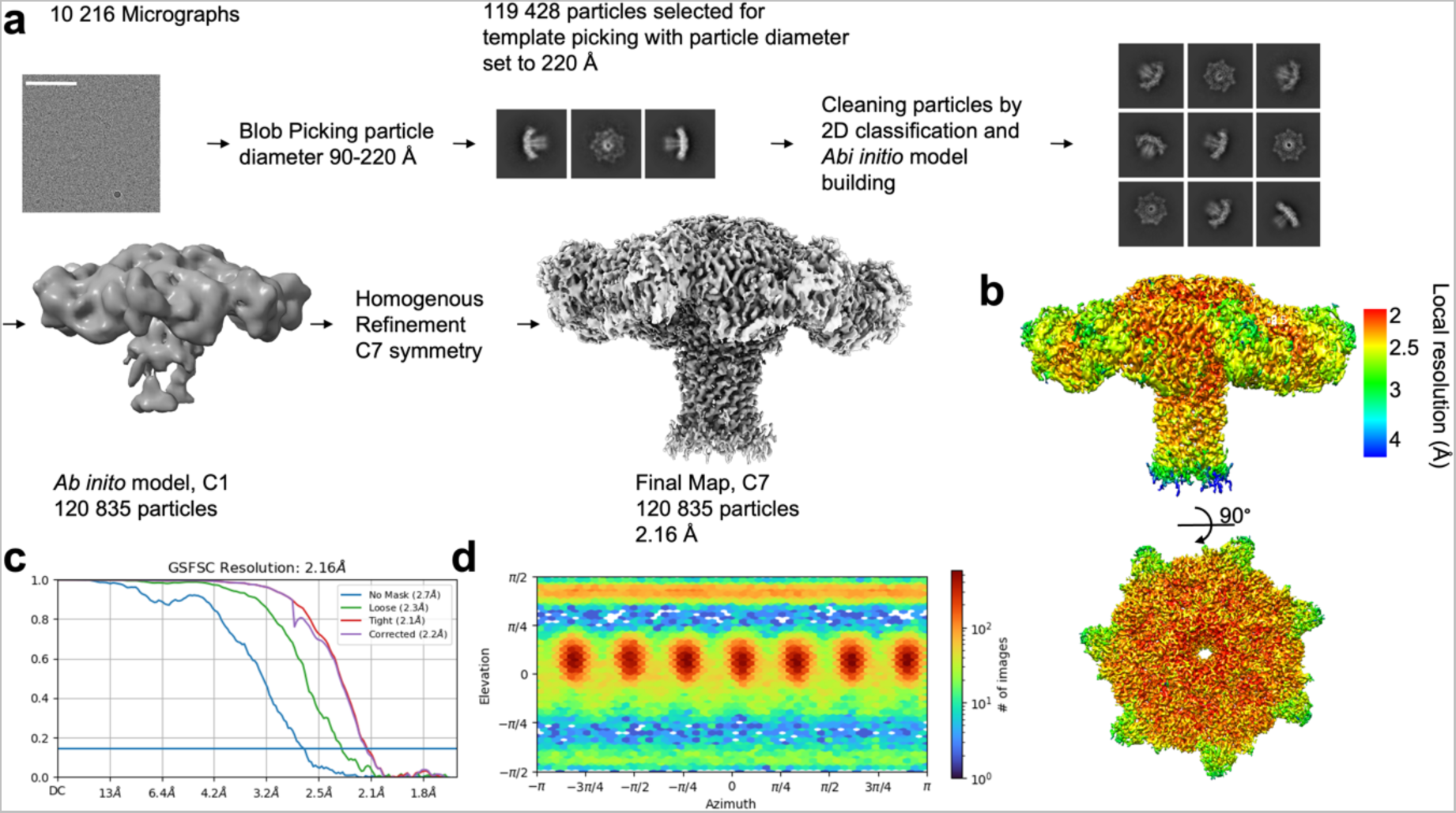
Cryo-EM processing of aerolysin mutant K238A in SMALP. **a** Flow chart of the data processing in cryoSPARC. Scale bar represent 100 nm. b Final 3D reconstruction of aerolysin K238A in SMALP colored by resolution. c Golden-standard Fourier shell correlation curves for the 3D reconstruction. d Angular distribution of the particles included in the final reconstruction.

**Supplementary Figure 1c:**
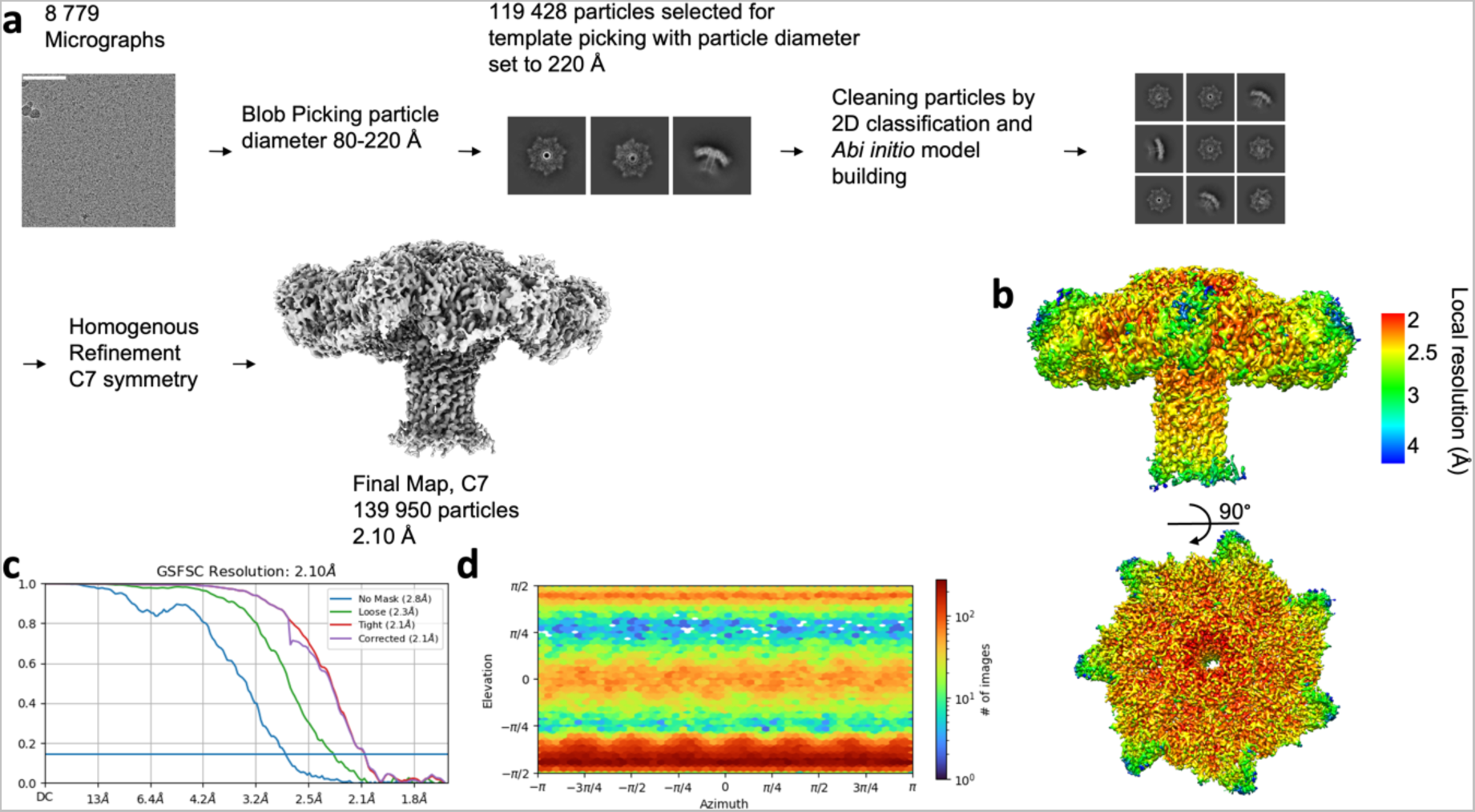
Cryo-EM processing of aerolysin mutant K238A/K244A in SMALP. **a** Flow chart of the data processing in cryoSPARC. Scale bar represent 100 nm**. b** Final 3D reconstruction of aerolysin K238A/K244A in SMALP colored by resolution. **c** Golden-standard Fourier shell correlation curves for the 3D reconstruction. **d** Angular distribution of the particles included in the final reconstruction.

**Supplementary Figure 1d:**
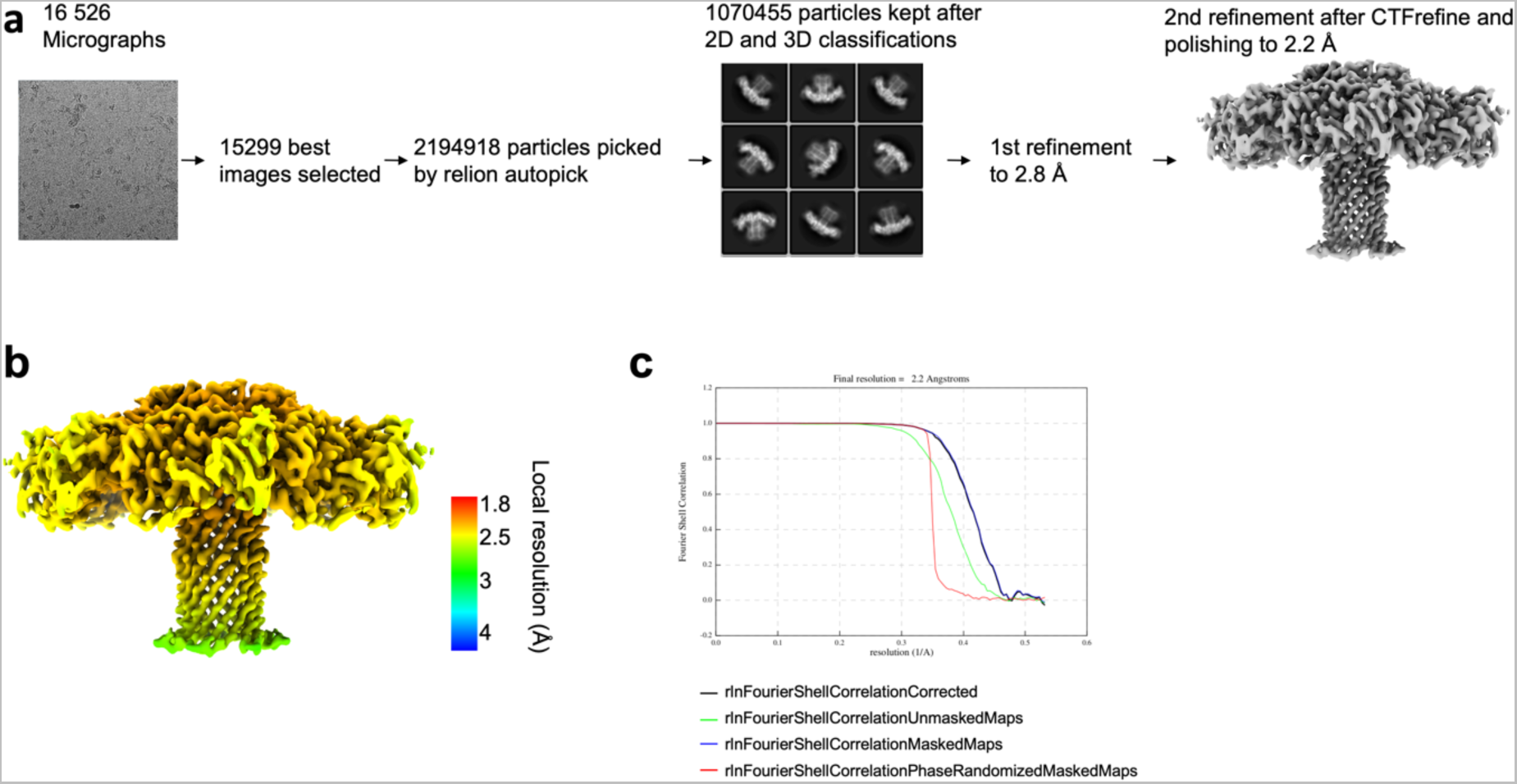
Cryo-EM processing of aerolysin WT in amphipol. **a** Flow chart of the data processing in Relion. Scale bar represent 100 nm**. b** Final map colored by resolution. **c** Golden-standard Fourier shell correlation curves for the 3D reconstruction.

**Supplementary Figure 1e:**
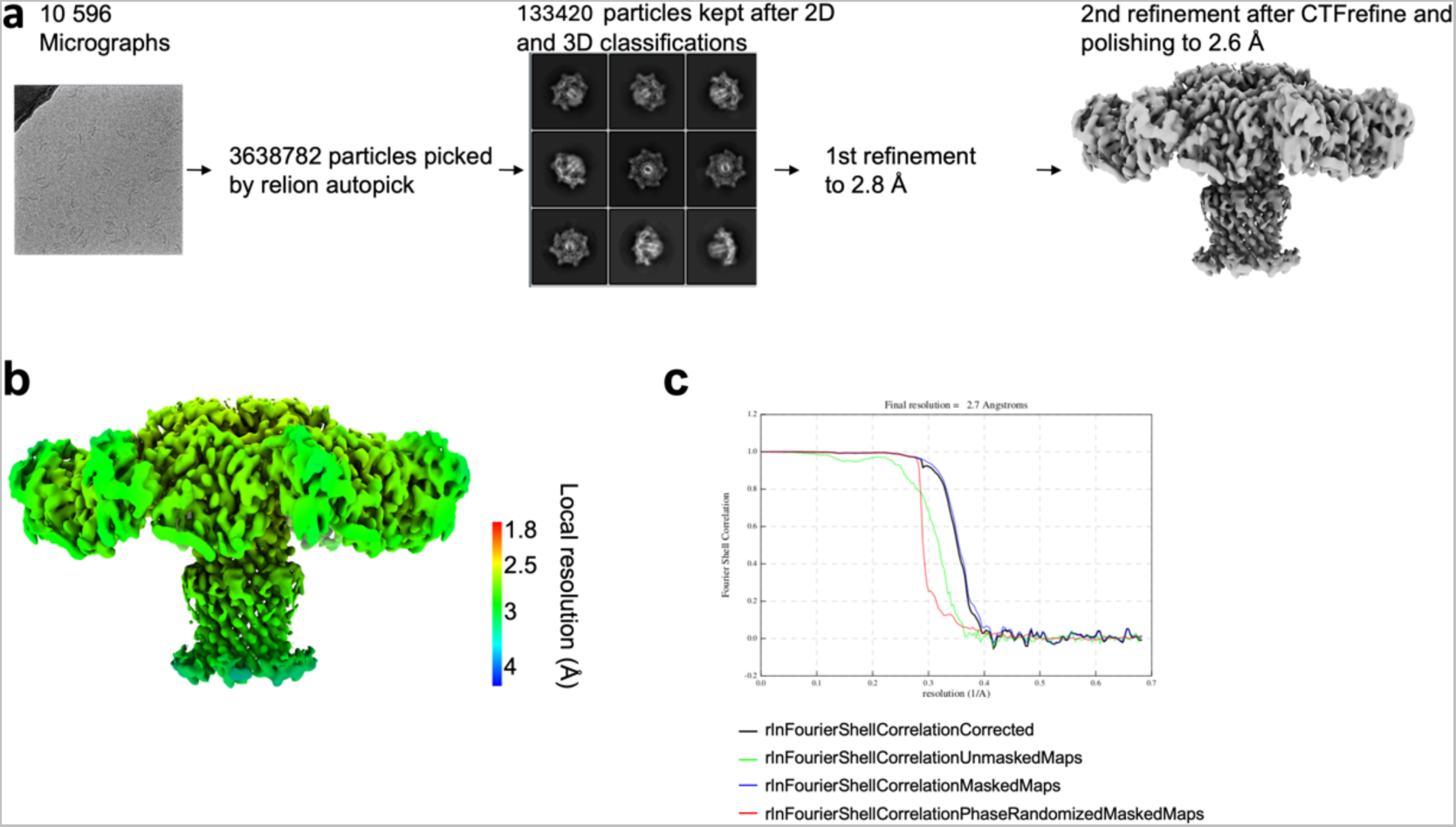
Cryo-EM processing of aerolysin WT in LMNG:CHS. **a** Flow chart of the data processing in Relion. Scale bar represent 100 nm. **b** Final map colored by resolution. **c** Golden-standard Fourier shell correlation curves for the 3D reconstruction.

**Supplementary Figure 1f:**
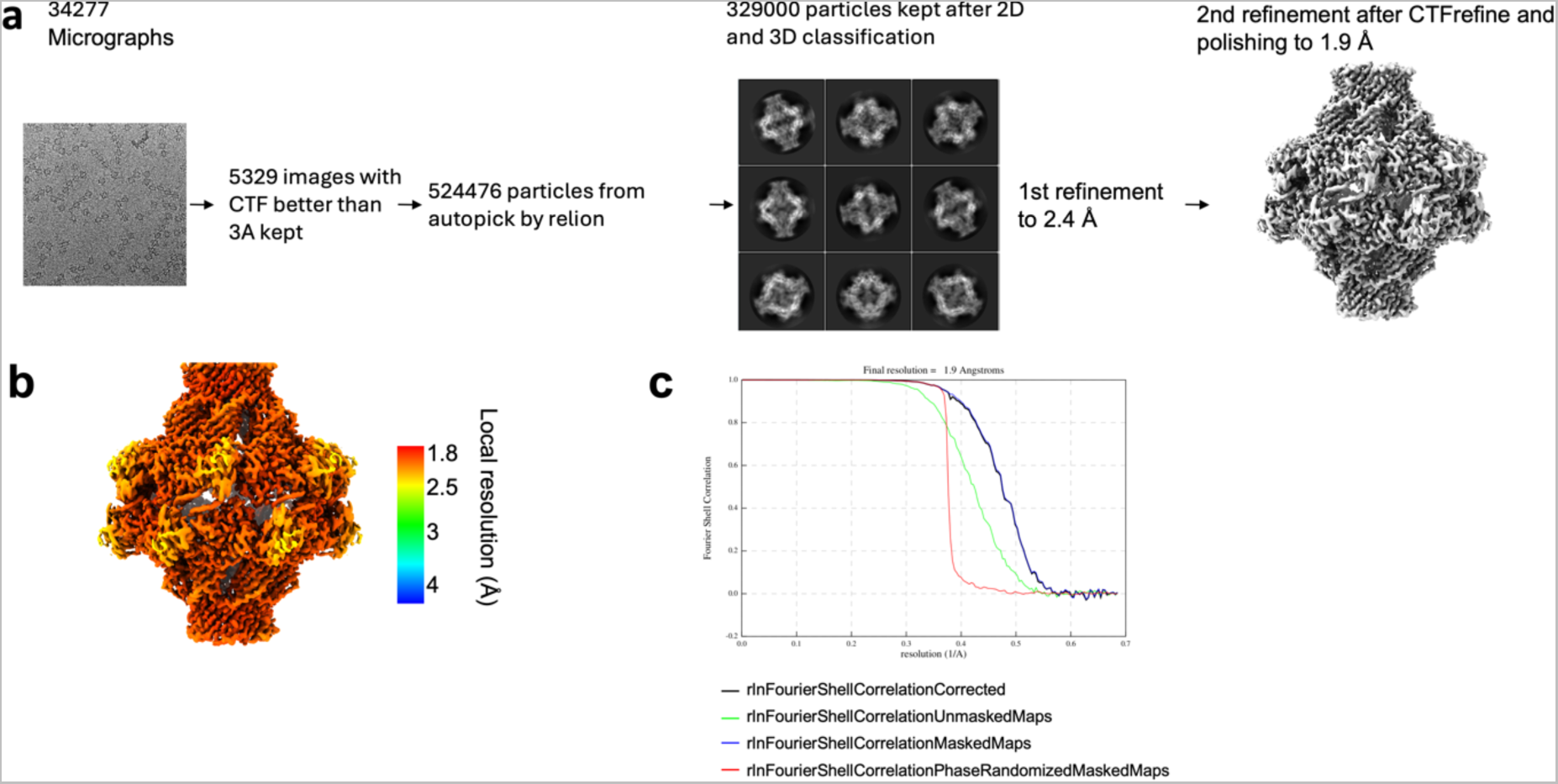
Cryo-EM processing of aerolysin mutant Y221G. **a** Flow chart of the data processing in Relion. Scale bar represent 100 nm. **b** Final map colored by resolution. **c** Golden-standard Fourier shell correlation curves for the 3D reconstruction.

**Supplementary Figure 2:**
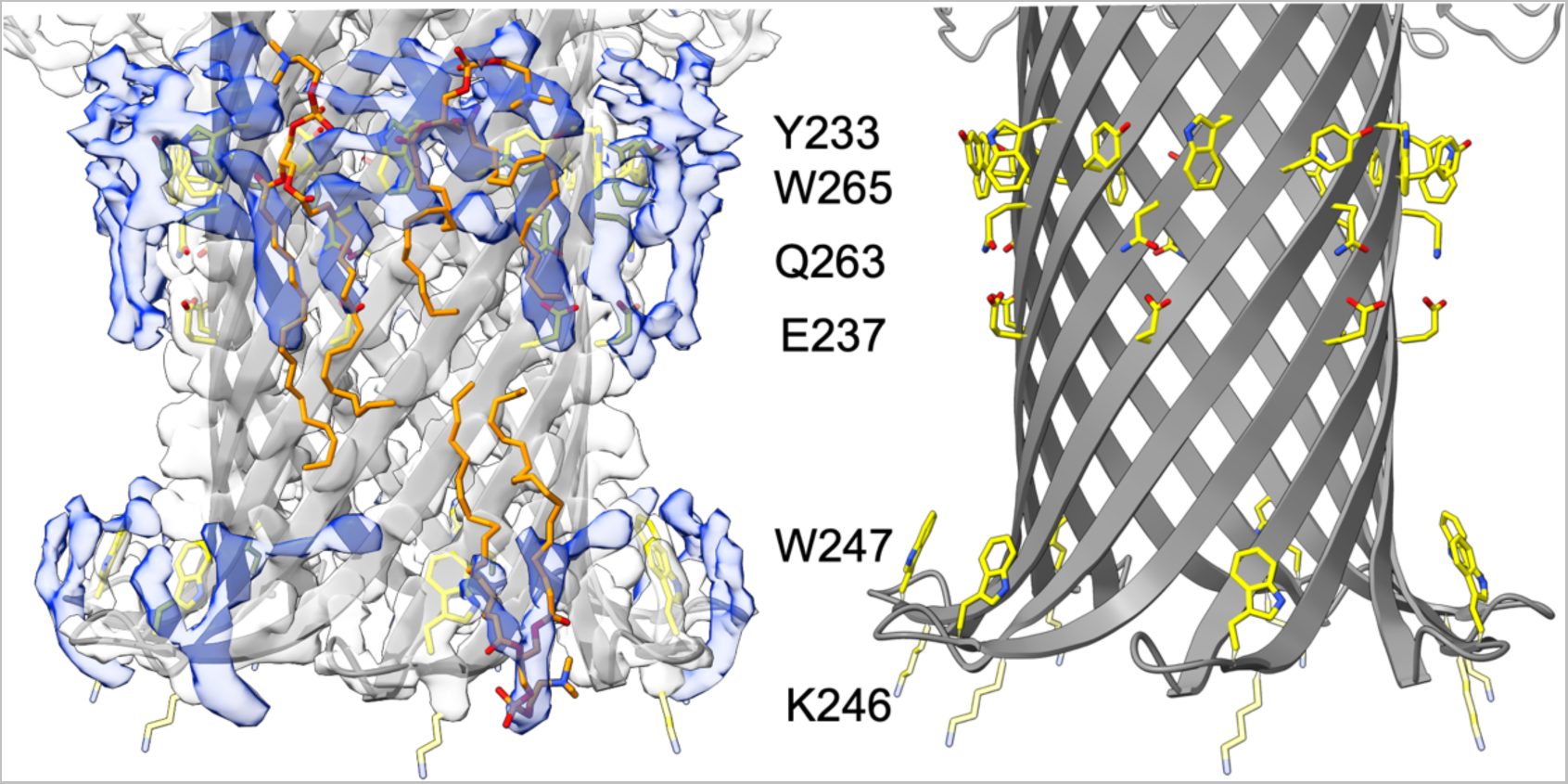
Docking of DOPC into extra density of the aerolysin in SMALP cryoEM structure. Zoom into the barrel region with the membrane lining residues highlighted in yellow, and the extra density in blue. Docked lipids are shown in orange. On the left the structure without the density map is shown.

**Supplementary Figure 3:**
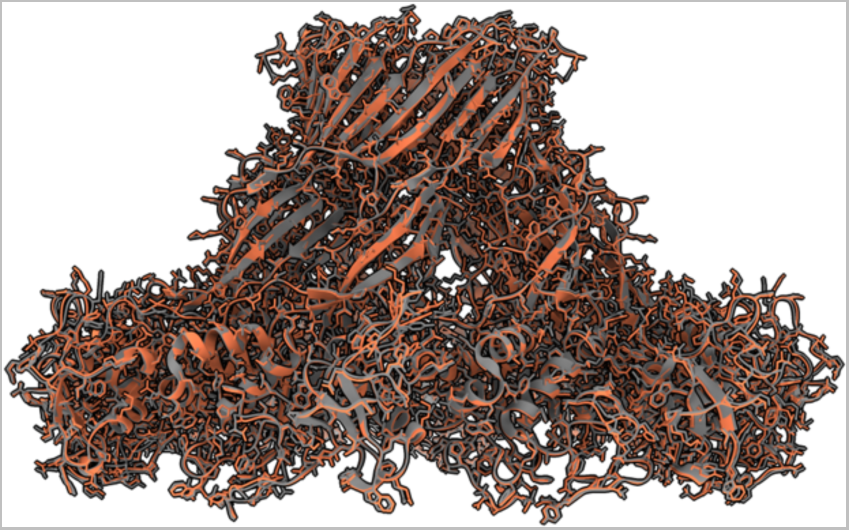
Aerolysin mutant Y221G trapped in its prepore state. In orange the new high-resolution structure is shown. In gray the previous PDB structure (5JZH).

**Supplementary Figure 4:**
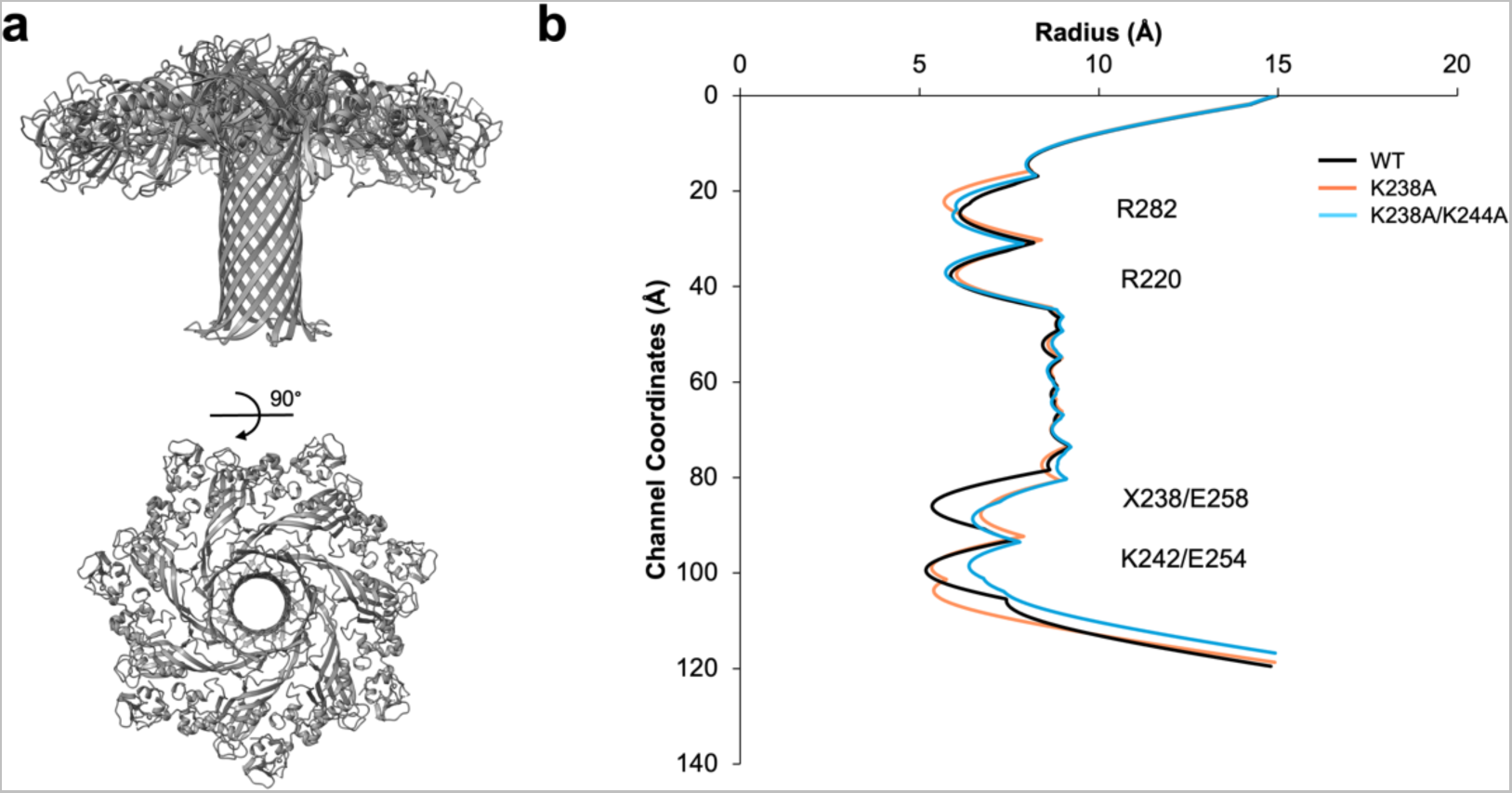
Aerolysin Mutant K238A/K244A. **a** Cryo-EM structure of the aerolysin mutant K238A/K244A in SMALP from the top and the side. **b** Comparison of the pore radii for the aerolysin wt (black), K238A mutant (orange) and the K238A/K244A mutant (blue).

**Supplementary Table 3:** CryoEM maps processing and model refinement statistics.

| Protein | WT SMALP | K238A SMALP | K238A K244A SMALP | WT LMNG | Aerolysin Y221G | Aerolysin WT in Amphipol |
| --- | --- | --- | --- | --- | --- | --- |
| **Access code** |  |  |  |  |  |  |
| PDB | 9FM6 | 9FNP | 9FNQ |  | 9FMX | 9FML |
| EMDB | EMD-50549 | EMD-50601 | EMD-50602 | EMD-50578 | EMD-50576 | EMD-50562 |
| **Data collection and processing** |  |  |  |  |  |  |
| Microscope | TFS Titan Krios G4 | TFS Titan Krios G4 | TFS Titan Krios G4 | TFS Titan Krios G5 | TFS Titan Krios G5 | TFS Titan Krios G5 |
| Detector | Falcon IV | Falcon IV | Falcon IV | Falcon IV | Falcon IV | Falcon IV |
| Recording mode | electron-counting mode (EER) | electron-counting mode (EER) | electron-counting mode (EER) | electron-counting mode (EER) | electron-counting mode (EER) | electron-counting mode (EER) |
| Magnification | 96000 | 96000 | 96000 | 165000 | 165000 | 130000 |
| Voltage (kV) | 300 | 300 | 300 | 300 | 300 | 300 |
| Total dose (e<sup>-</sup>/Å<sup>2</sup>) | 50 | 50 | 50 | 40 | 40 | 40 |
| Nominal under focus range (μm) | 0.8 - 1.7 | 0.8 - 1.7 | 0.8 - 1.7 | 0.8 - 2.5 | 0.8 - 2.5 | 0.8 - 2.5 |
| Pixel size (Å) | 0.83 | 0.83 | 0.83 | 0.73 | 0.73 | 0.94 |
| Number of movie micrographs | 10825 | 10216 | 8779 | 10596 | 34277 | 16526 |
| Number of molecular projection images in map | 89384 | 120835 | 139950 | 133420 | 329000 | 1070455 |
| Symmetry | C7 | C7 | C7 | C7 | D7 | C7 |
| Map resolution (Å) | 2.17 | 2.16 | 2.1 | 2.6 | 1.9 | 2.2 |
| Map sharpening B-factor | 47.6 | 55 | 38.8 | -65.8 | -35.32 | -43.55 |
| **Model refinement and validation** |  |  |  |  |  |  |
| Residues | 2884 | 2793 | 2870 |  | 5936 | 2954 |
| RMSD BondLength (4σ) | 0.009 | 0.006 | 0.003 |  | 0.005 | 0.003 |
| RMSD Bond Angles (4σ) | 1.153 | 0.878 | 0.567 |  | 0.634 | 0.656 |
| Ramachandran Outliers (%) | 0 | 0 | 0 |  | 0 | 0.24 |
| Ramachandran Allowed (%) | 1.47 | 1.78 | 0.99 |  | 2.95 | 3.33 |
| Ramachandran Favored (%) | 98.53 | 98.22 | 99.01 |  | 97.05 | 96.43 |
| Rotamer outliers (%) | 0.29 | 1.19 | 1.16 |  | 1.13 | 1.69 |
| Clash score | 0.61 | 1.76 | 2.25 |  | 3.61 | 7.32 |
| Molprobity score | 0.7 | 0.99 | 1.05 |  | 1.36 | 1.81 |
| EMRinger score | 6.6 | 6.4 | 5.56 |  | 6.55 | 6.08 |

